# Dairy wastewater grease stabilizes *in situ* mesophilic biomethanation for H_2_-to-CH_4_ conversion

**DOI:** 10.64898/2026.06.09.731101

**Authors:** Maria L. Ruiz-Lorenzo, Angela Lao-Zea, Antonio D. Moreno, Federico Ferrari, Israel Diaz, Javier Contreras, Raquel Iglesias, Silvia Suarez, Miguel G. Acedos

## Abstract

Power-to-Gas technologies are emerging as a key strategy to integrate surplus renewable electricity into energy systems, through the conversion of green hydrogen into methane. However, the practical implementation of biological *in situ* biomethanation is still constrained by operational and design requirements that are incompatible with most existing anaerobic digestion infrastructures. This study demonstrates a stable and efficient mesophilic (37°C) *in situ* biomethanation process driven by substrate-induced microbial selection rather than relying on continuous hydrogen supply. Anaerobic digesters co-digesting sewage sludge from a wastewater treatment plant with lipid-rich greases recovered from dairy wastewater developed a pre-adapted hydrogenotrophic consortium capable of effective CO_2_-H_2_ conversion under mesophilic conditions. Long-term operation confirmed the robustness and persistence of this microbial structure. Upon H_2_ addition, methane concentrations up to 82 % were achieved under atmospheric pressure, without biogas recirculation, with hydrogen-to-methane conversion efficiencies up to 90% and methane productivities of 1.64 NL_CH4_.L^−1^d^−1^. 16SrRNA-based microbial community analysis revealed that dairy grease co-digestion selectively enriched hydrogenotrophic methanogens, particularly *Methanospirillum*, together with syntrophic fatty-acid-degrading bacteria such as *Syntrophomonas*, promoting efficient interspecies hydrogen transfer. Importantly, the lipid co-substrate enabled the establishment and long-term stability of the hydrogenotrophic pathway independently of hydrogen availability, mitigating challenges associated with intermittent renewable energy supply. Overall, these findings challenge the common reliance on thermophilic conditions, continuous hydrogen input, pressurization, and gas recirculation in *in situ* biomethanation, demonstrating that substrate-driven microbial selection can replace conventional engineering requirements such as thermophilic operation or reactor modifications, providing a simpler and scalable strategy for mesophilic *in situ* biomethanation.

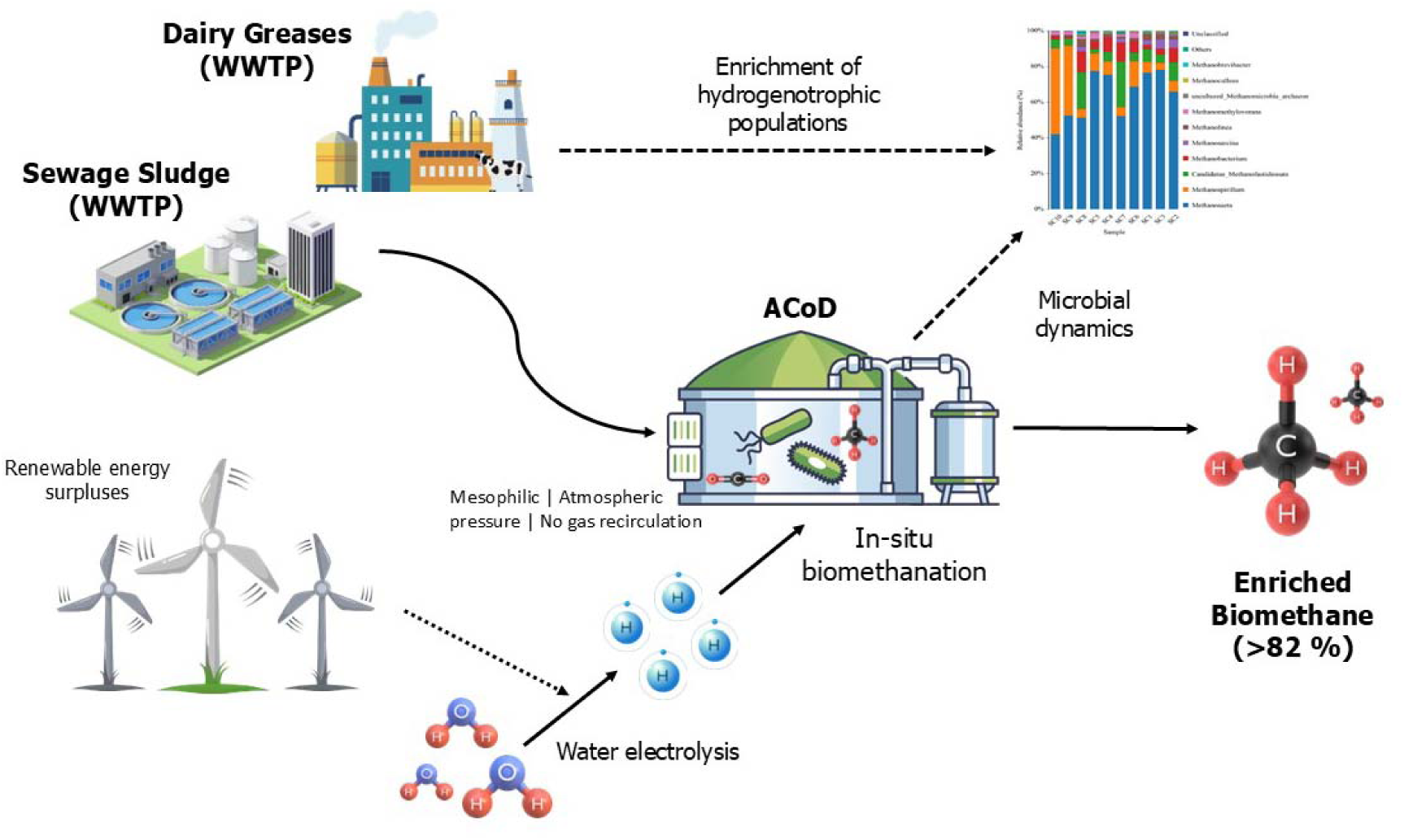
Graphical Abstract.

**Highlights:**

– Lipid-assisted co-digestion promotes stable biogas and biomethane production
– Dairy wastewater greases enable mesophilic in situ biomethanation
– An enriched hydrogenotrophic methanogenic consortium yields >82% CH_4_
– 70–90% H_2_-to-CH_4_ conversion efficiency under mesophilic, unpressurized conditions
– Substrate-driven microbial selection enables in situ biomethanation in WWTP digesters

## 1. Introduction

Today’s society faces major energy and environmental challenges, including rising global energy demand and increasing greenhouse gas emissions due to high consumption of fossil fuels. As a result, interest in anaerobic digestion (AD) as a sustainable technology for generating biogas and biomethane from organic waste has grown in recent years. AD not only contributes to energy production and security but also meets waste management objectives by reducing organic pollutants and greenhouse gases (GHG) [1]. This biomethane also contributes to energy security by reducing dependence on imports of fossil-based natural gas. The biogas produced by the anaerobic digestion of organic waste is mainly composed of methane (CH_4_) and carbon dioxide (CO_2_). CO_2_ can be transformed into biomethane, obtaining a high-purity renewable fuel suitable for injection into natural gas networks or direct use as vehicle fuel [2]. This process of transforming CO_2_ into CH_4_ is known as methanation and can be carried out through chemical catalysis or biological processes.

The integration of biological methanation technologies has gained increasing attention when compared to methanation by chemical catalysis because it allows for a more complete valorisation of the biogas stream [3]. An emerging strategy for biogas upgrading is biomethanation with green hydrogen or power-to-gas technologies. By relying on renewable hydrogen, typically produced by water electrolysis, biomethanation enables the transformation of the CO_2_ fraction of biogas into additional CH_4_, enhancing the overall energy recovery from organic residues. This process, known as hydrogen-mediated biomethanation, can follow two main approaches: in situ and ex situ biomethanation. In the in situ configuration, hydrogen is injected directly into the anaerobic digester, allowing the conversion of both endogenous CO_2_ and the exogenously supplied electron donor within the same reactor [4]. This approach has the advantage of not requiring additional reactors or separation units, making it potentially easier to integrate into existing infrastructure. However, its operational behaviour depends strongly on the microbial community dynamics, the characteristics of the substrate being digested, and the stability of the process under fluctuating hydrogen supply. Within this Power-to-Gas framework, co-substrates that simultaneously enhance methane productivity and mitigate the physicochemical disturbances associated with H_2_ addition are particularly attractive for retrofitting existing anaerobic digesters. In *in situ* biomethanation, substrate selection is especially relevant because the same reactor must support both conventional anaerobic digestion and biological H_2_ /CO_2_ conversion, making co-substrates a potential operational tool for process stabilization [3–4].

*Ex situ* biomethanation systems have been considered better conversion scenarios when compared to *in situ* strategies in power-to-gas technologies. This is due to the better control of H_2_ and CO_2_ injection, which creates favourable and stable conditions for the growth and maintenance of hydrogenotrophic methanogenic populations. This approach promotes hydrogenotrophic methanogen activity and enables biomethane purities above 96%, suitable for network injection [3]. In contrast, intermittent H_2_ injection during *in situ* biomethanation can impair process performance, as sustained hydrogen availability is often required to enrich and maintain hydrogenotrophic methanogens; under mesophilic conditions, this may lead to reduced hydrogenotrophic activity due to competition with alternative pathways and insufficient microbial adaptation [5–7].

Regarding in situ technology, the number of studies remains limited, and to date, most have not succeeded in directly producing high-purity biomethane [8–9]. Several studies have investigated *in situ* biomethanation under mesophilic conditions, highlighting both the potential and the limitations of this approach. For instance, Lebranchu et al. (2019) evaluated a pilot-scale anaerobic digester (100□L) equipped with a dense membrane for hydrogen injection, demonstrating that methane production increased proportionally with hydrogen supply; however, the process was found to be limited by CO□ availability and mass transfer phenomena, leading to residual hydrogen at higher injection rates [10]. Similarly, Alfaro et al. (2019) explored hydrogenotrophic methanation using membrane-based reactor configurations, reporting high hydrogen conversion efficiencies but under optimized conditions that enhance gas–liquid transfer and microbial selection [8]. These studies highlight that achieving efficient hydrogen conversion in mesophilic systems often relies on engineered reactor configurations and controlled operational conditions. Nevertheless, integration of in situ biomethanation into conventional digesters is attractive because it enables simultaneous waste stabilisation and CO_2_ conversion without new reactor construction and it has recently been demonstrated that it is a scalable technology [11–12]. However, in situ biomethanation still faces several critical limitations. One major challenge is the increase in pH typically observed when hydrogen is introduced. This pH rise results from the consumption of dissolved CO_2_ and bicarbonate, which serve as key components of the natural buffering system in anaerobic digesters. From this perspective, acidic co-substrates such as dairy greases are particularly interesting because their low pH can partially counterbalance the alkalinisation caused by CO_2_ consumption during hydrogenotrophic methanogenesis [13–15]. Another limitation, and probably the mainly bottle neck of in situ systems, is the low solubility of hydrogen which leads to inefficient mass transfer and requires intense mixing to maintain homogeneous conditions [3]. Although acidic dairy greases do not directly overcome H_2_ gas-liquid transfer limitations, their high methane potential may improve the overall energy recovery of the system, making *in situ* biomethanation more attractive under the moderate H_2_ loading rates compatible with conventional digesters [16]. Acetate accumulation is another recurrent issue in in situ systems. When hydrogen and CO_2_ are present, homoacetogens can outcompete hydrogenotrophic methanogens under certain conditions, redirecting H_2_ towards acetate production instead of methane [17]. In this context, co-substrates that promote syntrophic metabolism are of special interest, since lipid degradation is typically coupled to syntrophic fatty-acid oxidation and may help maintain the microbial interactions required to redirect reducing equivalents towards methane rather than acetate accumulation [17–18]. There are also thermodynamic and microbial considerations: hydrogenotrophic methanogens are typically more robust under thermophilic conditions, meaning that mesophilic reactors often struggle to maintain stable hydrogenotrophic populations [8]. This need is particularly relevant under mesophilic conditions, where hydrogenotrophic methanogens generally show slower enrichment and lower robustness than under thermophilic operation, thereby increasing the value of co-substrates able to favour their establishment without increasing temperature [19]. The experiments run at 55 °C have reported methane yields above 0.22 NL_CH□_NL_H□_^□1^, although higher values may reflect contributions from substrate degradation [20]. However, some researchers have demonstrated that bioaugmentation strategies using hydrogenotrophic archaea can be implemented to improve yields under mesophilic conditions, achieving CH_4_ production rates similar to those obtained under thermophilic conditions close to 1.0 L_CH□_.L□¹.d□¹ using whey as a substrate [19]. The need to shift digesters from mesophilic to thermophilic conditions represents an additional barrier, as this transition requires significant modifications in heating systems, control strategies, and operational procedures [12, 20–21]. Recent large-scale studies (>1000 m³) have shown that low or intermittent hydrogen dosing can still achieve high conversion rates [11, 22]. Despite these advances, the overall implementation of *in situ* biomethanation still requires solutions that ensure stable mesophilic operation, particularly with respect to maintaining balanced microbial communities, which are often more robust under thermophilic conditions, as well as mitigating pH fluctuations. A key knowledge gap therefore persists during *in situ* biomethanation: how to achieve stable mesophilic operation under continuous hydrogen supply without process inhibition or community destabilisation.

One promising *in situ* biomethanation strategy relies on the co-digestion of sewage sludge from WWTP (one of the most common substrates used in AD) with acidic organic co-substrates. This approach counteracts the alkalinisation caused by hydrogen injection, helping to maintain stable reactor conditions [23]. Sewage sludge is one of the most common substrates used in full-scale anaerobic digesters due to its continuous availability in municipal wastewater treatment plants. However, sewage sludge usually exhibits relatively low biodegradability, low C/N ratio, and high nitrogen and inorganic content. Conversely, dairy industries generate large amounts of wastewaters and by-products, such as whey, dairy greases, and treatment sludges, which are rich in lipids, proteins, and volatile solids and have a high methane potential [14–15]. These residues are challenging to treat in conventional WWTP systems because of their high organic load and biodegradability but are excellent candidates for co-digestion. In the European Union alone, approximately 400 billion litres of whey-rich wastewater are generated annually [24–25]. Dairy greases, which accumulate in separators and pre-treatment units, are particularly rich in lipids and therefore contribute to higher CH_4_ yields, although they may induce long-chain fatty acid (LCFAs) inhibition if fed individually [26]. Their acidic nature makes them ideal for buffering pH increases during in situ biomethanation, helping maintain stable reactor conditions. Co-digestion with these residues also improves the nutrient balance, enhances buffering capacity, and can mitigate other challenges such as LCFA accumulation, ammonia inhibition, or deficiencies in trace elements [27–28].

The introduction of hydrogen into anaerobic digesters not only influences reactor chemistry but also has a profound impact on the microbial community structure. H_2_ availability exerts a strong selective pressure favouring hydrogenotrophic methanogens, which directly utilise H_2_ and CO_2_ to produce CH_4_. Genera such as *Methanoculleus*, *Methanothermobacter*, *Methanospirillum*, and *Methanosarcina* commonly dominate in hydrogen-amended systems [29]. These organisms belong to the phylum Euryarchaeota and are considered robust markers of stable biomethanation. At the same time, acetoclastic methanogens typically decline under high H_2_ availability, while homoacetogenic and syntrophic bacteria may increase in abundance [30]. Homoacetogens compete directly with hydrogenotrophs for H_2_, often consuming up to 40% of the available H_2_ under stress conditions, although even in well-performing reactors around 5% of H_2_ may be diverted towards acetate production [31]. Substrate composition may also modulate this competition, as lipid-rich co-substrates can select for syntrophic fatty-acid degraders and hydrogen-scavenging methanogens, thereby potentially giving the hydrogenotrophic pathway a competitive advantage under mesophilic conditions [31–32]. Maintaining syntrophic interactions between VFAs-oxidising bacteria and hydrogenotrophic archaea is therefore essential to avoid propionate or butyrate accumulation, which can lead to instability and process inhibition [18]. Additionally, hydrolytic and fermentative microorganisms, particularly those belonging to Bacteroidetes, Proteobacteria and Firmicutes, play essential roles in the degradation of complex substrates such as lipids, proteins, and carbohydrates, especially when co-digesting dairy residues. Their activity is crucial for ensuring efficient hydrolysis and fermentation, enabling subsequent methanogenesis [32]. Numerous studies have focused on studying hydrogenotrophic microbial populations in mesophilic and thermophilic conditions, and even in the transitions between both temperatures, and they all agree that hydrogenotrophic populations are stable and more efficient at thermophilic temperatures [33–35]. Palú et al. achieved stable in situ biomethanation under mesophilic conditions using bioaugmentation strategies with *Methanoculleus bourgensis*, and increased the biomethane yield of the mesophilic process by 11% [19]. Understanding how these microbial populations respond to H_2_ addition and co-digestion strategies is vital to develop stable operational regimes for *in situ* biomethanation.

Taken together, these considerations make acidic dairy greases an attractive co-substrate for mesophilic in situ biomethanation. First, their acidic character can help offset the pH increase caused by CO_2_ consumption during hydrogenotrophic methanogenesis. Second, their high lipid content provides an additional methane-rich substrate that can improve the overall energy recovery of the process. Third, lipid degradation is strongly associated with syntrophic microbial networks, which may favour the establishment of hydrogen-scavenging methanogens and thereby enhance the stability of H_2_/CO_2_ conversion under mesophilic conditions. These combined features suggest that substrate selection may represent a simpler alternative to more intensive engineering solutions such as thermophilic operation, pressurization, or gas recirculation. Importantly, this approach shifts biomethanation from a reactor-engineering challenge to a biologically-driven process design strategy [13–14, 18, 26].

This study investigates continuous co-digestion of sewage sludge from a WWTP with acidic dairy greases from dairy WWTP, combined with in situ biomethanation using H□. Dairy greases were selected to buffer the pH increase resulting from CO□ consumption during biomethanation. The results demonstrate that these lipid-rich co-substrates stabilize hydrogenotrophic archaeal populations under mesophilic conditions, enabling high methane production rates (up to 1.64□L_CH□_□L□¹□d□¹). Importantly, while in situ biomethanation has been successfully demonstrated under mesophilic conditions [16], several studies report limitations related to slower hydrogenotrophic methanogen growth rates and competition with alternative metabolic pathways, which can affect process efficiency and stability; in contrast, thermophilic conditions are generally associated with enhanced hydrogenotrophic activity and more robust process performance [36–37]. By overcoming these constraints without requiring thermophilic operation, this study provides a simple and scalable strategy to enhance methanogenic activity, addressing a key microbiological bottleneck and supporting the practical, cost-effective implementation of in situ biomethanation at industrial scale.

## 2. Materials and methods

### 2.1. Substrates and inoculum

Dairy grease (Supplementary Figure S1) as collected from the WWTP of a cheese production facility owned by Entrepinares S.A.U. in Fuenlabrada (Madrid, Spain), while mixed sewage sludge (SS) was obtained from the combined primary and secondary treatment decanted sludge of a municipal WWTP in Madrid. The cheese factory produces 15 tonnes of cheese whey per week. The waste used in this study did not require pre-treatment. An 80:20 w:w co-digestion mixture of sewage sludge and cheese factory fats was used according to Lao-Zea. et al 2026 [38]. The physicochemical characterization of all individual substrates and co-digestion blends is summarized in **Table 1**. The inoculum was a digestate from a mesophilic biogas plant located at the same WWTP (Madrid). This plant processes sewage sludge with crude glycerol as a co-substrate. This WWTP digester is fed with sewage sludge at a hydraulic retention time (HRT) of 30-35 days. Upon lab arrival, the inoculum was screened to remove debris and degassed at 37□°C for 5 days. The inoculum characteristics are also detailed in **Table 1**.

**Table 1.** Results of the physicochemical characterisation of waste, co-digestion mixtures and inoculum. Values are expressed as the mean ± standard deviation of three independent replicates (n = 3).

| Parameter | Inoculum | SS-WWTP | DG | SL-WWTP + DG (80:20) |
| --- | --- | --- | --- | --- |
| pH | 7.2 ± 0.5 | 6.3 ± 0.4 | 5.4 ± 0.1 | 5.9 ± 0.2 |
| τCOD (g.Kg<sup>-1</sup>) | 30.1 ± 10.2 | 42.1 ± 13.1 | 225.8 ± 33.9 | 90.6 ± 42.6 |
| $s$COD (g.Kg<sup>-1</sup>) | 1.3 ± 0.6 | 1.2 ± 0.8 | 41.8 ± 3.0 | 10.0 ± 0.2 |
| TS (%) | 3.1 ± 0.1 | 4.1 ± 0.6 | 19.1 ± 0.3 | 6.2 ± 1.5 |
| VS (%) | 2.0 ± 0.1 | 3.0 ± 0.4 | 17.2 ± 0.7 | 5.0 ± 1.3 |
| H (%) | 96.9 ± 0.2 | 95.9 ± 0.6 | 81.0 ± 0.3 | 93.8 ± 1.5 |
| Ash (%) | 1.1 ± 0.1 | 1.1 ± 0.7 | 1.8 ± 0.8 | 1.20 ± 0.6 |
| C<sub>T</sub> (%bs) | 31.1 ± 0.7 | 37.6 ± 1.4 | 59.5 ± 2.2 | 49.4 ± 0.5 |
| N<sub>T</sub> (%dw) | 4.5 ± 0.2 | 5.3 ± 0.3 | 3.7 ± 0.2 | 5 ± 1 |
| S<sub>T</sub> (%dw) | 1.2 ± 0.12 | 0.9 ± 0.1 | 0.3 ± 0.03 | 1 ± 1 |
| VFAs (g.L<sup>-1</sup>) | < 0.01 | 0.9 ± 0.1 | 8.7 ± 1.2 | 3.7 ± 0.7 |
| Acetic acid (g.L<sup>-1</sup>) | n.d | 0.5 ± 0.6 | 1.6 ± 0.4 | 1.1 ± 0.1 |
| Propionic acid (g.L<sup>-1</sup>) | n.d | 0.2 ± 0.3 | 3.7 ± 0.4 | 1.4 ± 0.4 |
| Butyric acid (g.L<sup>-1</sup>) | n.d | 0.1 ± 0.1 | 1.9 ± 0.4 | 0.6 ± 0.1 |

n.d: not detected; SL-WWTP: sewage sludge; DG: dairy greases

### 2.2. Continuous anaerobic digestion (AD) process

3-L continuously stirred tank bioreactors (CSTRs) were used in this study. In addition, a 20-L stainless steel CSTR was employed to carry out the *in situ* biomethanation experiments with H_2_ (Supplementary Figure S2). The reactors designated as R1, R2, and R3 in this study have a volume of 3 L, whereas the reactor designated as R4 has a volume of 20 L. The reactors were equipped with a thermostatic water jacket to maintain the operating temperature, and mixing was provided by a Rushton-type impeller driven by an external motor (50 rpm). The digesters were automatically fed and discharged daily using programmable peristaltic pumps. The operating temperature was controlled at 37 °C (mesophilic conditions). The organic loading rate (OLR) and hydraulic retention time (HRT) were adjusted throughout the semi-continuous experiments. The feeding strategy and corresponding OLR applied in each experimental phase are summarized in Table 2. Biogas production was measured using MilliGascounter devices (Ritter, Germany), and the accumulated biogas was collected in Tedlar bags.

**Table 2.**
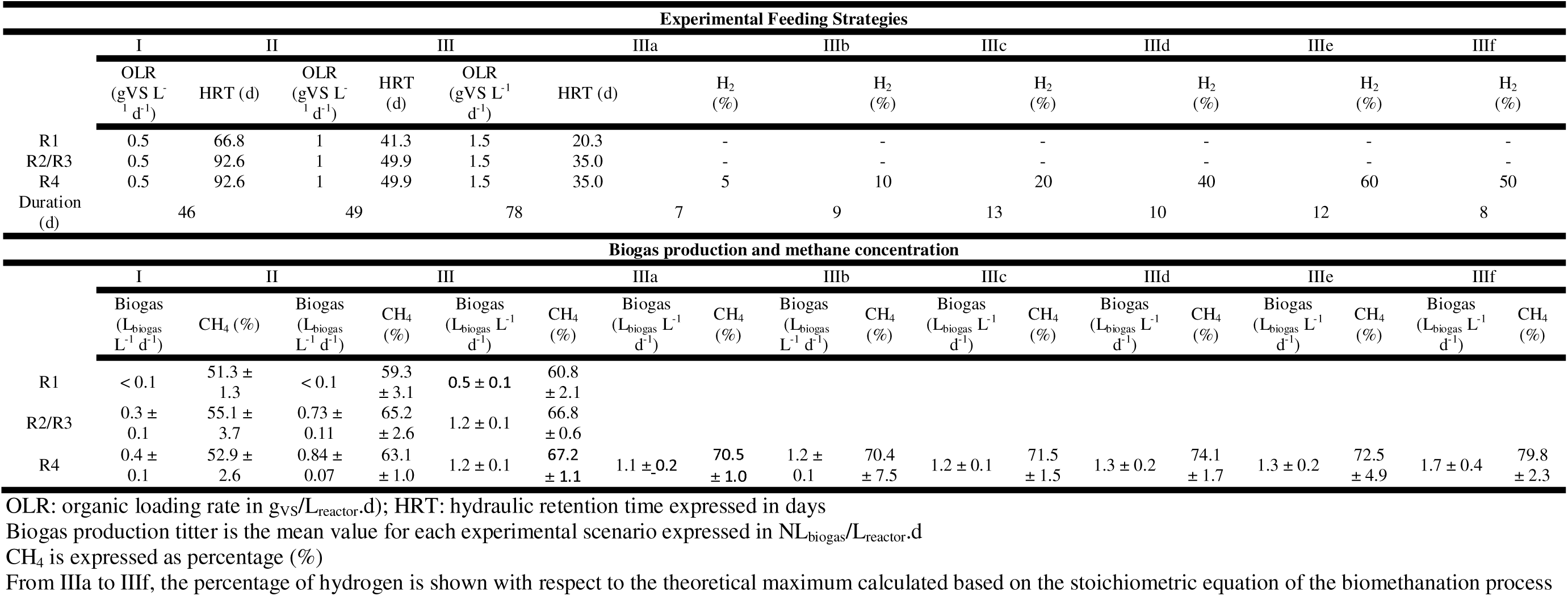
Feeding strategy and biogas and biomethane production titers for reactors R1 (single digestion sewage sludge), R2 and R3 (replicas for co-digestion of sewage sludge and cheese factory fats 80:20 w/w), reactor R4 (sewage sludge and cheese factory fats 80:20 w/w in which hydrogen has been injected)

### 2.3. H_2_ generation and feed

A hydrogen generator (HG ST BASIC 120) with a maximum production rate of 120 mL/min and a purity of 99.99 % was used (Supplementary Figure S2). The system operates at a maximum pressure of 10 bar and is based on proton exchange membrane (PEM) technology using distilled water, allowing continuous operation for up to 24 hours. The PEM technology produces high-purity H_2_ (typically 99.99% or higher) and enables rapid adjustment of H_2_ output, making these electrolysers suitable for applications coupled with intermittent renewable energy sources. Hydrogen was fed into the digester using a mass flow controller (Alicat Scientific, USA); the flow rate of hydrogen added was adjusted daily based on the flow rate of biogas produced. To this end, a percentage was calculated based on the stoichiometric ideal (4 moles of H_2_ for every mole of CO_2_) and was gradually increased until we approached that ideal value. The gas was injected into the digester via a porous plate 10 cm in diameter at the base of the digester (pore size 0.5 mm).

### 2.4. Analytical methods

#### 2.4.1. Physicochemical analysis for substrates, inoculum and digestates in anaerobic digestion tests

A comprehensive physicochemical characterization was performed on the substrates, co-digestion mixtures, inoculum, and digestates collected during the experiments. The parameters measured included total solids (TS), volatile solids (VS) (see FigS6-S9), pH (see FigS6-S9), FOS/TAC ratio (see FigS6-S9), total and soluble chemical oxygen demand (COD_T_ and COD_S_), total carbon (TC), total organic carbon (TOC), total inorganic carbon (TIC), total Kjeldahl nitrogen (TKN), ammonium nitrogen (NH_4_–N), orthophosphate (PO_4_–P), total phosphorus (TP), alkalinity, and concentrations of selected micronutrients. Standard analytical procedures were applied for TS and VS determination [39]. TS were quantified by drying the samples at 105□±□2□°C until a constant mass was reached, while VS were assessed by incinerating the dried material at 550□°C for six hours using a muffle furnace. TC and TIC were measured using a Primacs ATC100-IC Analyzer (Skalar, Netherlands); TOC was calculated by difference. The pH was measured directly using a 913 pH-meter (Metrohm, Switzerland). TKN, which includes both organic nitrogen and ammonia, was determined using the Kjeldahl method with an integrated digestion and distillation system comprising the K-439 Automatic Digester, K-415 Scrubber, and K-365 distillation unit (Buchi, Switzerland). COD_T_ and COD_s_ were analysed using pre-prepared HANNA test tubes (HI 93754C-25, range: 0–15000□mg L^−1^). The samples were subjected to thermal digestion at 150□°C for a period of two hours using a HANNA HI 839800 reactor. Following digestion, absorbance was measured with a HANNA HI 83214 photometer to quantify COD levels.

Volatile fatty acids (VFAs) were identified and quantified via high-performance liquid chromatography (HPLC). The analyses were carried out on a Waters Alliance 2695 system (Waters, MA, USA) equipped with a 410 refractive index detector. Separation was achieved using an Aminex HPX-87H column (Bio-Rad Laboratories, Hercules, CA) maintained at 65□°C, with 5□mM H_2_SO_4_ as the mobile phase at a flow rate of 0.6□mL/min.

#### 2.4.2. Microbial community analysis

Microbial population analyses were carried out to determine the effect of the addition of grease and H_2_. DNA was extracted at the beginning and at the end of the digestion using the TGuide S95 Magnetic Universal DNA kit (DP812, TiangenBiotech, Beijing, China). The concentration of the extracted DNA was assessed using the Nanodrop2000 (Thermo Fisher Scientific, Wilmington, USA). Specific primers were designed to target on full-length amplicon (Supplementary Table S1). Target sequences were amplified by PCR and its products were purified, quantified and homogenized to generate SMRT bell library. The qualified library was sequenced on the PacBio Revio (Pacific Biosciences, Menlo Park, USA). Library construction and sequencing were performed at Biomarker Technologies (BMKGENE) GmbH. After amplification, the PCR products are analysed by LabChip GX (Perkin Elmer, Waltham, USA) for fragment analysis and assessment of the integrity. Library QC was performed on the libraries, and the qualified ones were processed for sequencing on PacBio Sequel II platform. Pacbio Sequel II initially generated files in BAM (Binary Alignment Map) format, which were further converted into CCS files by SMRT (Single Molecule Real Time) link. Resulting sequences were analysed for taxonomic and functional composition.

#### 2.4.3. Gas analysis

The CH_4_, CO_2_, H_2_ sulphide, oxygen and nitrogen concentration in the produced biogas was determined daily by two methods. Firstly, we use for a daily measurement an MRU Biogas Optima 7 gas analyser. Also, for biogas composition, samples were analysed using a gas chromatograph (8860 GC, Agilent Technologies) equipped with FID and TCD detectors. For each measurement, 0.5 mL of biogas were manually injected in triplicate to ensure analytical reproducibility. The equipment was fitted with two PLOT Q columns for the separation of CO_2_ and light hydrocarbons and one Molsieve column for permanent gases (H_2_, O_2_, N_2_ and CH_4_), enabling accurate quantification of the main biogas components throughout the semi-continuous trials.

## 3. Results and Discussion

### 3.1. Substrates and inoculum characterization

**Table 1** lists the physicochemical characterization of inoculum, sewage sludge, dairy greases and co-digestion mixture. TS and VS ranged from 4.1-19.1 % and 3.0 % to 17.2 %, respectively, consistent with those reported for similar wastes in the bibliography [15, 27]. Likewise, both COD_T_ and COD_S_, are also similar with those found in the literature [15, 27]. The pH values of the substrates are within the expected range from 5.4 to 6.3, with dairy greases being notably more acidic than sewage sludge. The concentration of VFAs was highly variable, ranging from 0.9 g.L^−1^ in the Ss to 8.7 g.L^−1^ in dairy waste grease. Elevated VFAs, particularly in lipid-rich substrates, can inhibit methanogenic activity if not buffered appropriately [40–41]. Regarding carbon concentration, co-digestion can enhance microbial activity by balancing nutrient availability; however, none of the co-digestion mixtures achieved the optimal C/N ratio required for efficient biogas and biomethane production [15, 42–44]. Nevertheless, co-digestion mixtures containing sewage sludge can reduce total solids, thereby improving digester rheology and mass transfer. Among the substrates employed in this work, dairy waste grease stood out for its high biodegradability and energy potential, but its high VFAs content and acidity make it unsuitable for mono-digestion (Bella & Rao, 2022). Conversely, co-digestion with sewage sludge can supply the buffering capacity. Their combination, particularly at an 80:20 (w/w) ratio for SL-WWTP + Dairy Grease had previously demonstrated intermediate characteristics, in solids content, VFAs and COD, that could enhance the process stability while optimizing substrate utilization [38].

### 3.2. Stepwise OLR Experiment and *in situ* H_2_ driven biomethanation

The reactors were operated under a stepwise OLR increase strategy, as summarized in Table 2, to evaluate the effect of substrate composition, and subsequent H_2_ addition on process performance. A preliminary set of experiments was carried out to establish the necessary experimental protocol and operational conditions. The results are provided as supplementary material (Supplementary **Table S2**, Supplementary **Figure S3**).

Figure 1 shows the results for reactors R1–R3, which were fed with either sewage sludge alone (R1) or a co-digestion mixture of sewage sludge and dairy greases (80:20 w/w; R2 and R3), and R4, which evaluates the *in situ* biomethanation performance of the co-digestion mixture through controlled H_2_ injection. H_2_ feeding began at IIIa phase and was increased stepwise (5–60 % of stoichiometric CO_2_ demand) through phases a to f according to **Table 2**. As it can be seen in Figure 2, CH_4_ concentration in the produced gas increased in all co-digestion reactors compared with the sludge mono-digestion system, but the most significant enrichment was recorded in R4, where H_2_ was supplied. The enhanced performance is attributed to the high biodegradability of dairy greases, which promote syntrophic β-oxidation reactions releasing H_2_ and acetate—key intermediates for methanogenesis. These findings agree with previous studies showing that moderate fat co-substrate additions can increase methane yield by 20–40 % compared with sludge-only digestion, provided that LCFA_S_ inhibition is controlled [45–46].

**Figure 1.**
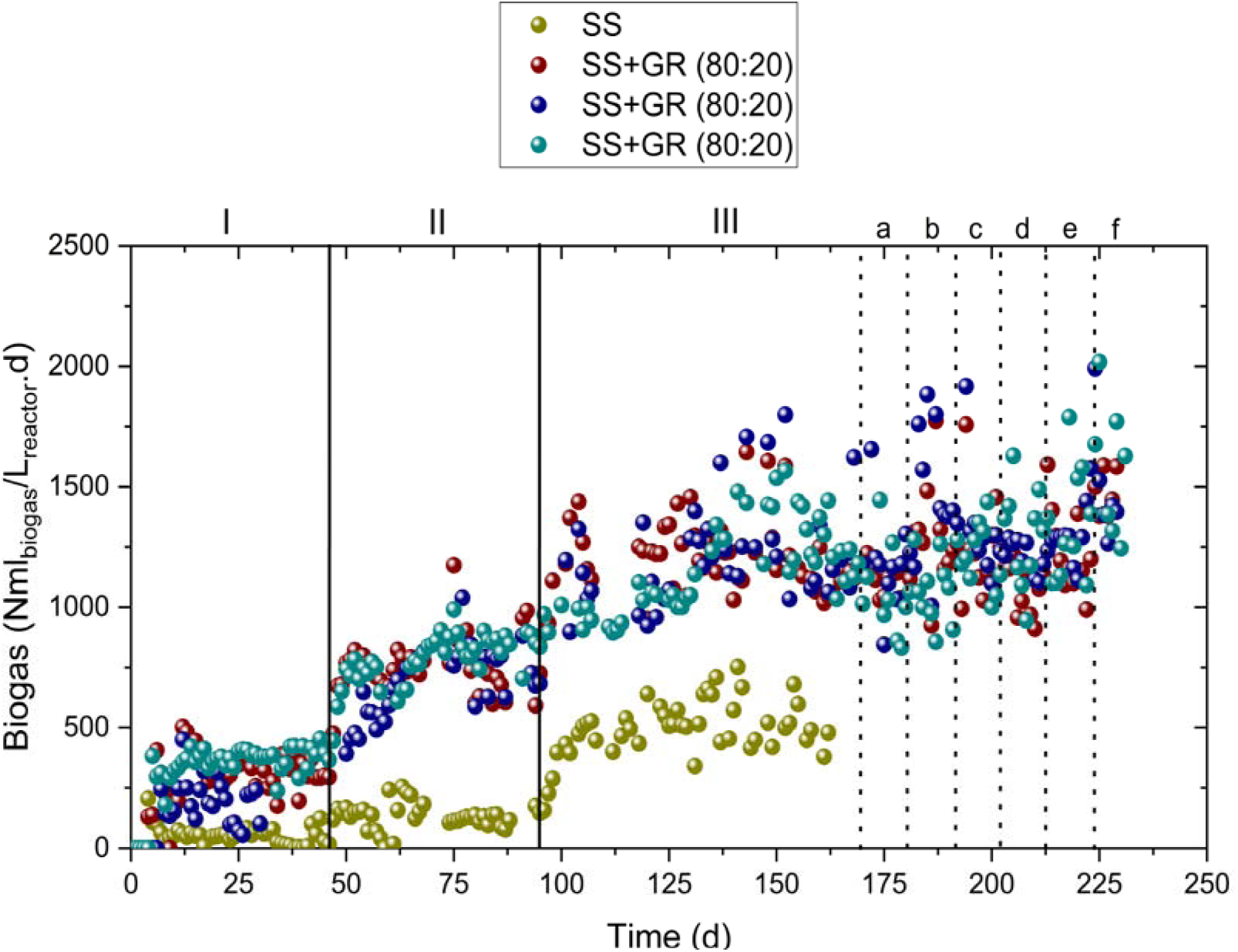
Time course of the mono-digestion of untreated sewage sludge (green; SS, R1) and co-digestion of untreated sewage sludge and dairy greases (80:20 w/w) (red and dark blue; SS+GR 80:20, R2 and R3), as well as the test of greases and sewage sludge with hydrogen addition (light blue, SS+GR, R4). The lines separate the different stages (IIIa-f) in which the changes in the feed rate of hydrogen to the digesters were carried out, as described in Table 2.

**Figure 2.**
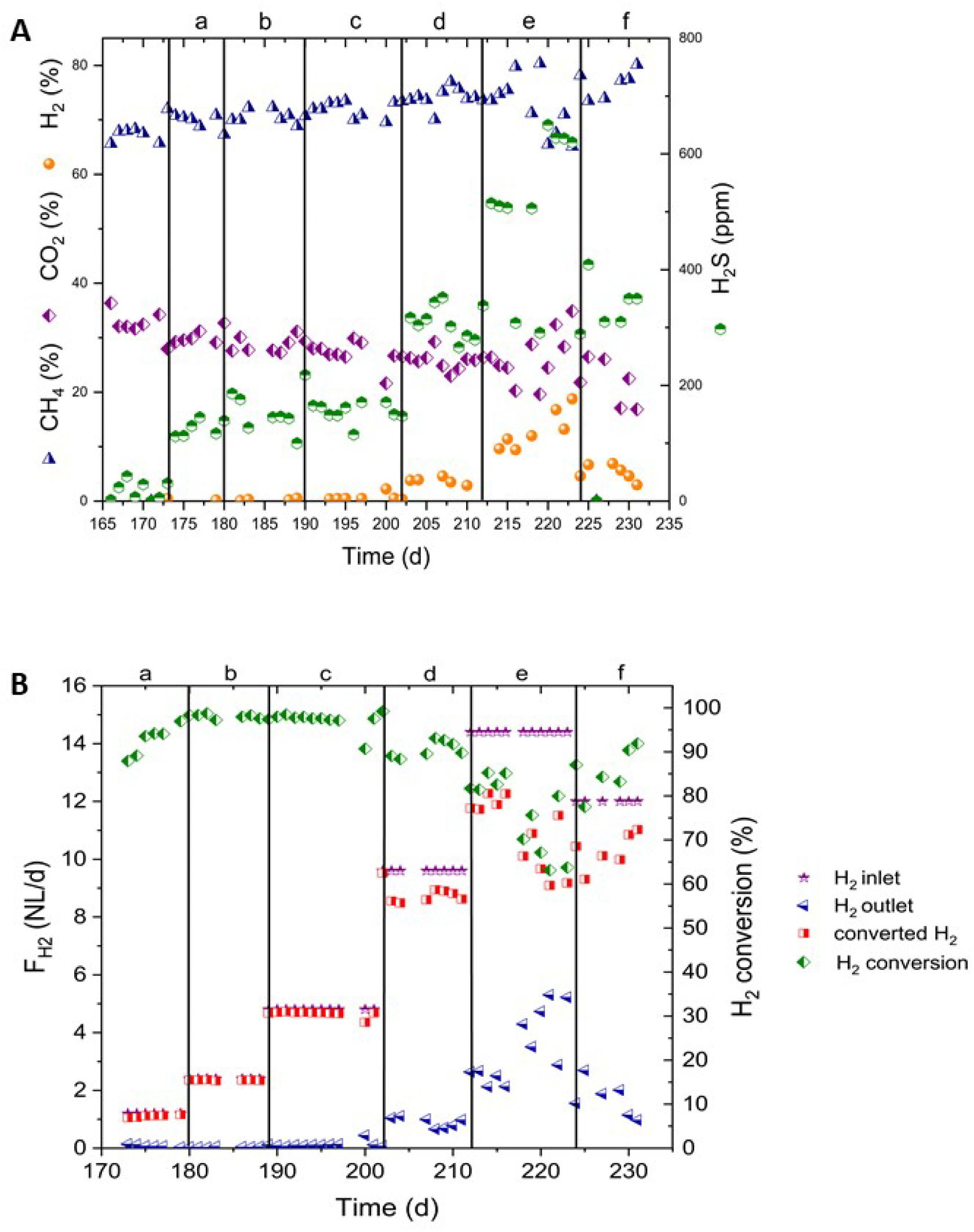
**A**) Biogas composition profile under biological methanation conditions in the R4 reactor (20L) following the progressive H_2_ injection strategy**; B)** Relationship between the H_2_ injection rate (input) and the percentage of H_2_ conversion according to output flows for each of the H_2_ addition stages. The different stages (a-f) in which the changes in the feed rate of hydrogen to the digesters were carried out are described in Table 2

Figure 2 illustrates the dynamic behaviour of this reactor during the hydrogen addition phases. Each increase in H_2_ flow led to a sharp increase in total methane flow rate, followed by a progressive stabilization of methane content up to 83 % CH_4_. This indicates that a large fraction of the injected H_2_ was effectively converted into CH_4_ rather than accumulating in the headspace. Importantly, this enrichment was achieved under mesophilic conditions (37□°C), which are often considered less favourable for hydrogenotrophic methanogens.

**Table 2** and Supplementary Table S3 list the biogas production and composition for each experimental stage. R1, operated in mono-digestion of sewage sludge from WWTP, exhibited a moderate increase in biogas production with increasing OLR, rising from 0.1 to 0.5 NL_biogas_.L_reactor_□¹ d□¹ between stages I and III, while CH□ concentration improved from 51.3 % to 60.8 %. In contrast, co-digestion in R2 and R3 substantially enhanced biogas production, leading to a more stable process performance. Biogas production increased from 0.26 to 1.19 NL_biogas_.L_reactor_□¹ d□¹ across the same OLR range, with CH_4_ contents consistently higher (55.1–66.8%). The addition of dairy greases greatly increased volumetric biogas productivity by approximately three to five-fold, when compared to mono-digestion at similar OLR, confirming the strong synergistic effect typically associated with lipid-rich co-substrates. Reactor R4, operated with the same co-digestion mixture, displayed similar behaviour to R2/R3 in stages without H_2_ addition. After H_2_ supplementation, a clear shift in gas composition was observed. While biogas volumetric productivities remained within comparable ranges (1.12--1.32 NL_biogas_.L_reactor_□¹ d□¹ for 5–50 % H_2_ addition), CH_4_ concentration increased progressively up to biomethane quality from 70.5 % to a maximum of 82 % at the highest H_2_ dosing stage (60 % H_2_). These values are consistent with those reported for in situ biomethanation systems, where methane contents in a similar range (75-81%) have been observed depending on reactor configuration and operating conditions [23, 46–47]. In addition, a significant increase was also noticed in the concentration of H_2_S when H_2_ is injected, rising from 2.8 ppm to 500 ppm. This phenomenon has been previously reported in the literature [23] and is associated with the dynamics of sulfur-transforming microbial communities under anaerobic conditions. In particular, sulfate-reducing bacteria (SRB) can utilize hydrogen as an electron donor, competing directly with hydrogenotrophic methanogens and leading to the reduction of sulfate to H_2_S. This competitive interaction has been widely described and can significantly affect both methane production and sulfide levels in anaerobic systems [48]. In the context of hydrogen addition, the increased availability of H_22_ may therefore stimulate SRB activity, resulting in elevated H_2_S concentrations. Additionally, changes in pH and carbonate equilibrium associated with CO_2_ consumption during biomethanation may further influence sulfide speciation and its transfer to the gas phase. These results highlight the importance of controlling sulfur cycling when implementing in situ biomethanation processes.

The efficient conversion observed suggests that the microbial community obtained with co-digestion of Ss and dairy greases was already metabolically prepared to utilize H_2_, likely due to the prior co-digestion with dairy greases. The β-oxidation of LCFAs by hydrolytic bacteria enriched hydrogenotrophic archaea (notably *Methanospirillum*), which could rapidly consume exogenous H_2_ when it became available [49]. LCFAs were degraded to acetate through the well-known LCFAs β-oxidation pathway and the combined β-oxidation and butyrate oxidation pathway [49]. This adaptive phenomenon has been described as a “biological preconditioning” effect, where lipid co-substrates favour the selection of microorganisms capable of sustaining syntrophic electron transfer and hydrogenotrophic methanogenesis [17, 30, 49]. No oscillations or acidification events were observed during this process, showing a stable CH_4_ fraction after each H_2_ pulse (Figure 2A). It is important to highlight that these results were obtained without the need of recirculating the biogas + H_2_ and clearly reflects the high buffering capacity of the system and the establishment of tight syntrophic coupling between hydrogen-producing and hydrogen-consuming guilds, thereby avoiding potential foaming issues and additional energy consumption commonly associated with gas recirculation strategies. It also highlights that *in situ* biomethanation can be operated intermittently—synchronized with renewable H_2_ availability—without losing microbial functionality, a key advantage for power-to-gas integration. It should be noted that efficient hydrogen conversion can also be achieved in mono-digestion systems without prior acclimation when low hydrogen loading rates are applied.

Figure 2B quantifies H_2_ utilization efficiency as a function of injection rate. Conversion remained high (70–90 %) up to approximately 40 % of the stoichiometric input, indicating efficient gas–liquid transfer and biological uptake. Despite this, a 10% content of unprocessed hydrogen is too high in terms of process costs, and future efforts must focus on developing processes in which there is no free hydrogen in the off-gas. At higher flow rates, conversion declined moderately, consistent with diffusion and solubility constraints inherent to atmospheric *in situ* systems [8, 22]. Nevertheless, no accumulation of H_2_ or process imbalance was detected, confirming that the microbial consortium could maintain metabolic homeostasis under variable redox conditions. During this period, CH_4_ production rates ranged from 0.45 L_CH□_.L_H2_^−1^ per day, when no H_2_ was injected, to 1.18 L_CH□_.L_H2_^−1^ per day, when 50 % of the stoichiometric H_2_ was supplied. This result is particularly noteworthy considering that our system operated under mesophilic conditions, whereas the literature values reported by M. Fachal-Suárez, S. Krishnan, S. Chaiprapat, D. González and D. Gabriel [20] (0.2–0.5 m³_CH_.m^−^³) were obtained under thermophilic conditions, which typically enhance CH_4_ production [20].

Overall, the two-step operational approach, a progressive OLR increase followed by a controlled H_2_ addition, demonstrated that mesophilic digesters co-fed with dairy greases can operate as bioactive methanation platforms. The pre-adaptation to lipidic substrates provided a stable microbial syntrophic network capable of immediate H_2_ assimilation, thus bridging the gap between conventional anaerobic digestion and power-to-gas biotechnology. This finding is particularly relevant for full-scale WWTP, where retrofitting existing mesophilic digesters for renewable H_2_ use represents a realistic route to achieve carbon-neutral operation. Although the resulting CH_4_ concentration does not reached sufficient quality for grid injection, it meets the requirements to be used as carrier for energy storage, enabling its direct use in combined heat and power (CHP) units within the same WWTP. This approach is especially relevant in light of recent EU objectives promoting energy independence and self-sufficiency of WWTPs, positioning biomethanation as a key enabling technology rather than solely a gas-upgrading pathway. However, under higher hydrogen loading conditions, process stability and conversion efficiency may be compromised due to mass transfer limitations and microbial competition. In this context, the use of co-digestion in the present study contributed to maintaining stable hydrogenotrophic activity and high methane production rates under more demanding operational conditions. These operational results further demonstrate that mesophilic digesters co-fed with dairy greases can maintain stable performance at organic loading rates typical of full-scale WWTP anaerobic digestion while efficiently converting externally supplied H_2_ into CH_4_. This functional robustness is directly linked to the underlying microbial community structure, which evolved toward lipid-degrading and hydrogenotrophic metabolisms. This finding demonstrates that process performance can be primarily governed by substrate-driven microbial selection, rather than by conventional process intensification strategies, redefining the role of co-substrates as active tools for reactor design in biomethanation systems.

### 3.3. Microbial community dynamics during co-digestion of sludge and dairy greases and hydrogen-assisted biomethanation

Figure 3 shows the relative abundance of archaeal phyla (Fig. 3A) and genera (Fig. 3B) obtained in the analyzed samples. The nomenclature of each sample can be observed in Table S4 in the Supplementary Material. Firstly, the most representative phyla in all samples are Halobacterota and Euryarchaeota. In the mono-digestion of sewage sludge, an increase in the abundance of Euryarchaeota and Thermoplasmatota is observed. In contrast, in the co-digestion with greases, Euryarchaeota shows a sustained increase over time, suggesting an enrichment of hydrogenotrophic methanogens and syntrophic associations dependent on H_2_ and methylamines. The phyla Thermoplasmatota and Crenarchaeota remain at low or relatively constant levels under all the evaluated conditions. Altogether, these results indicate that both the type of substrate and OLR employed modulate the composition of archaeal phyla, and that co-digestion with fats favors the proliferation of specific phyla associated with hydrogenotrophic methanogenesis.

**Figure 3.**
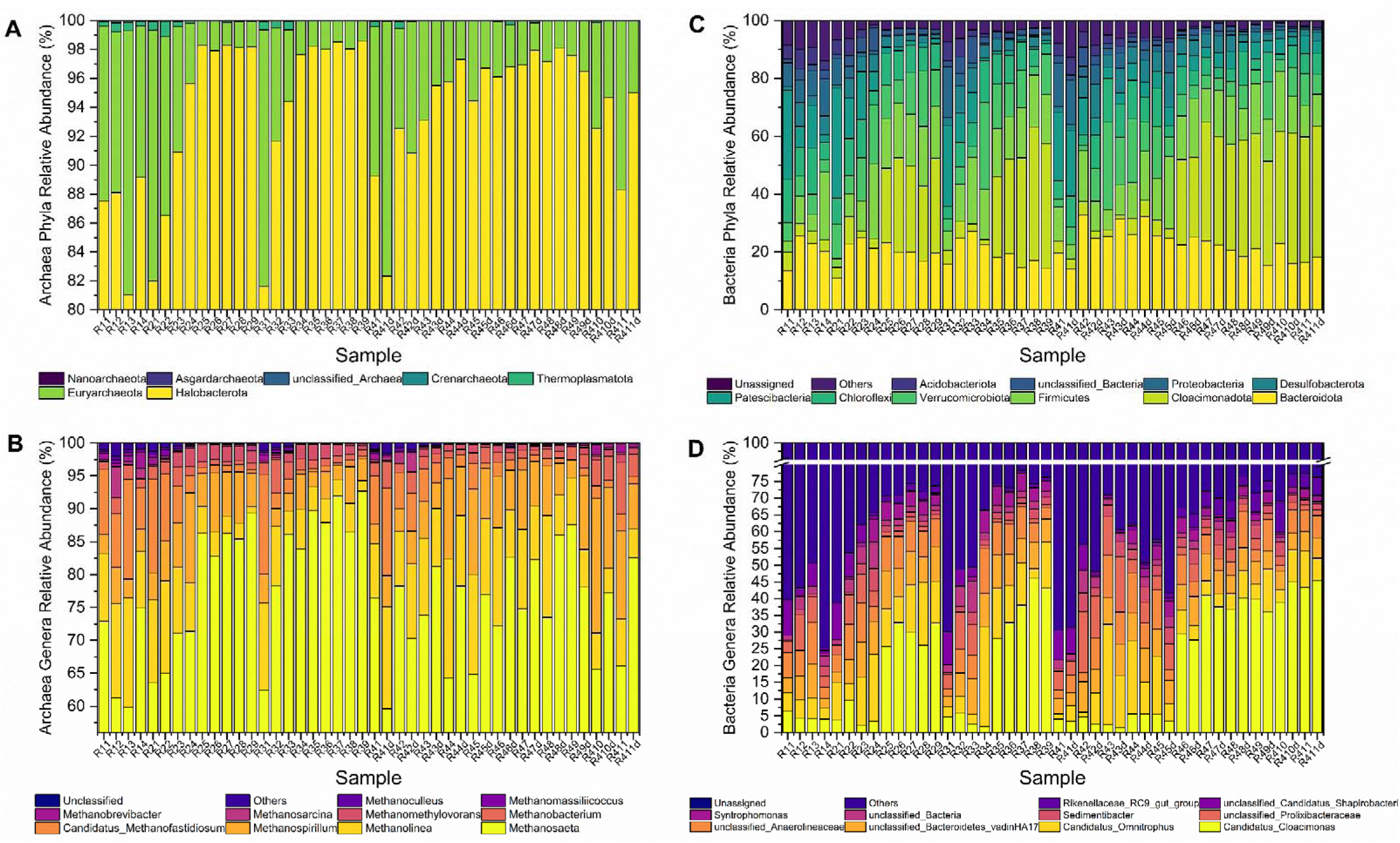
Histogram of relative abundance of main phyla (A) and genera (B) of archaea, and phyla (C) and 555 genera (D) of bacterial read.

Regarding archaeal genera (Fig. 3B), it should be noted that the genus *Methanosaeta* is the most abundant in all samples, but in this case, it can be said that the presence of fats in co-digestion favors the development of the genus *Methanosaeta* and *Methanospirillum* [50]. This difference is also clearly seen in Figure 4, which shows a comparative analysis of the abundance of the main archaeal genera under two different situations: monodigestion of sewage sludge vs. co-digestion of sewage sludge with cheese fats (Fig. 4A) and the effect before and after adding H_2_ (Pre and Post in Fig. 4B). These microorganisms have already been previously described in the literature, and their good capacity to produce CH_4_ when coupled with microorganisms that degrade LCFAs has been studied [46, 49].

**Figure 4.**
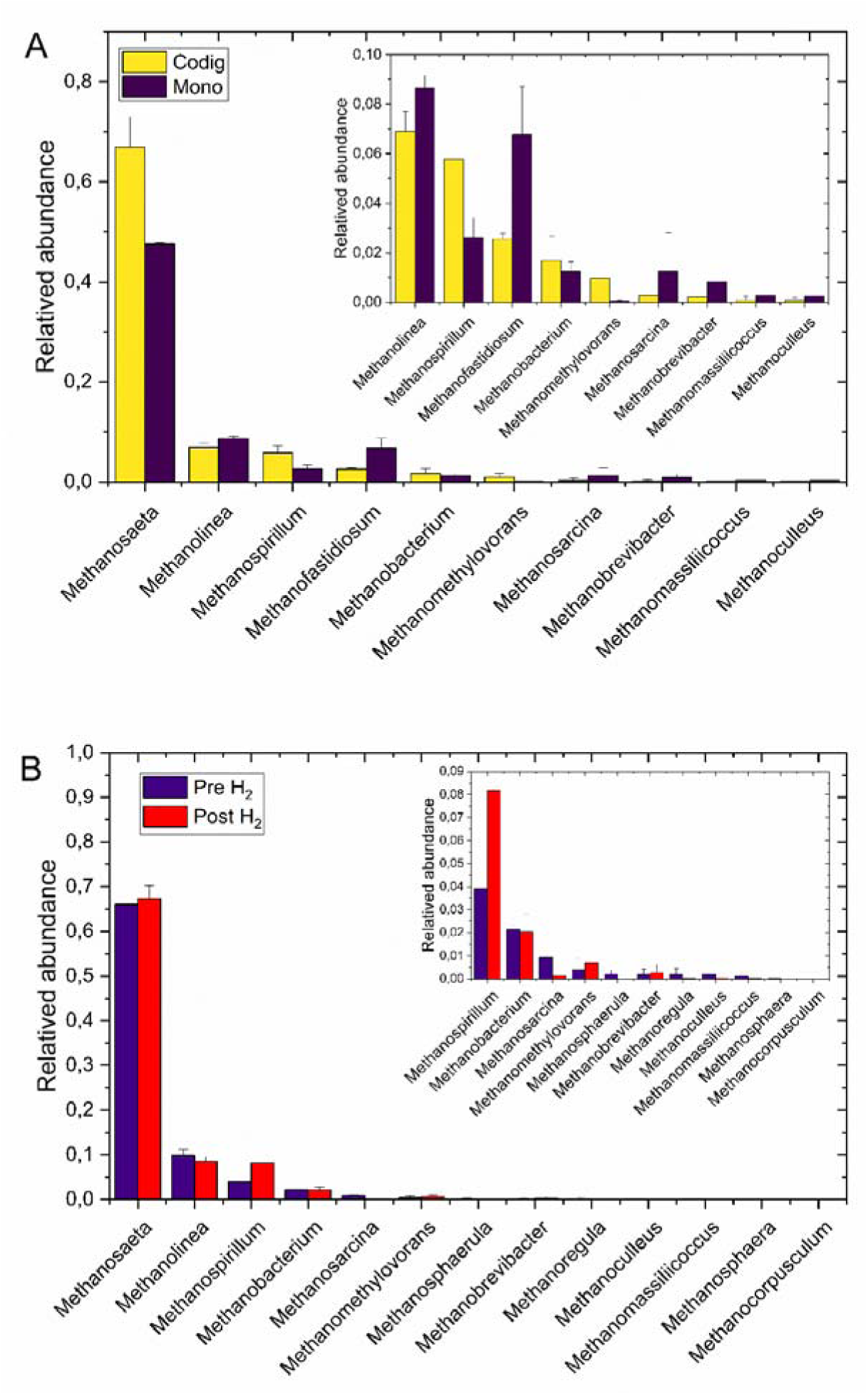
Comparative analysis of relative abundance of main archaea Genus comparing two different conditions. A) Mono-digestion samples and Co-digestion samples; B) Hydrogen suppling samples vs non-hydrogen suppling samples.

The reason why their abundance may have increased in these digesters is that the dairy fats (rich in long-chain fatty acids) are first hydrolyzed and fermented by hydrolytic bacteria capable of producing lipase enzymes in the digester, which subsequently release VFAs and, above all, H_2_ + CO_2_ during β-oxidation and acidogenesis. This excess H_2_ is the ideal substrate for *Methanospirillum*, which increases in abundance because it benefits from the high partial pressure of H_2_. Its increase indicates that the system is channeling a large part of the electron flow toward H_2_ (through lipid fermentation) and that methanogenesis is being directed more toward the CO_2_/H_2_ → CH_4_ pathway than toward the acetoclastic one.

On the other hand, sewage sludge has a high diversity of soluble substrates, including methylamines and nitrogen compounds from domestic and industrial wastewater [46, 51]. The mono-digestion of urban sludge generates a stable environment relatively rich in these compounds, which favors the enrichment of species related to the genus *Candidatus Methanofastidiosum*, as can be observed in the samples from mono-digestion sludge digesters (Figure 3B and Figure 4A).

Figure 3C and **3D** shows both the relative abundance of bacterial phyla and genera in the different samples, corresponding to digesters fed with sewage sludge in mono-digestion and digesters in co-digestion with dairy greases. In the Figure 3C it can be observed that in all samples the phyla Firmicutes, Bacteroidota, Chloroflexi, and Proteobacteria are present, which agrees with what has been reported in the literature about typical bacterial communities in anaerobic digestion [52–53]. However, the relative abundance of these groups varies depending on the type of substrate used. In digesters with co-digestion of dairy fats, an increase in Firmicutes can be seen, probably due to the ability of these phyla to degrade lipids and other polymeric molecules, generating VFAs and H_2_ that favor fermentation [45, 54]. On the contrary, Chloroflexi and Proteobacteria show a higher abundance in mono-digestion of sewage sludge, reflecting their role in the degradation of complex organic compounds and biomass diversity present in sludge [55]. Furthermore, the presence of dairy fats causes a notable increase in the abundance of Cloacimonadota and Verrucomicrobiota phyla associated with the fermentation of intermediates and the degradation of complex polysaccharides, respectively [56–57]. Moreover, in the Figure 5A shows how, when fats are co-digested, the medium is enriched with bacterial genera characteristic of lipid degradation, such as Syntrophomonas, *Bacteroidetes bacterium*, and Eubacterium WCHB1_07 was observed [49, 58–59]. These taxa are associated with fatty acid β-oxidation, H_2_ and formate production, and syntrophic associations with hydrogenotrophic archaea [59]. Their enrichment indicates a reinforced metabolic coupling between fermentative and methanogenic populations, promoting CO_2_ reduction to CH_4_ through hydrogenotrophic pathways. These results also indicate that the type of substrate modulates the structure of the bacterial community, favoring groups specialized in the degradation of predominant compounds and, therefore, directly influencing the metabolic dynamics of the digester, promoting the degradation of LCFAs, producing H_2_, and increasing CH_4_ yields.

**Figure 5.**
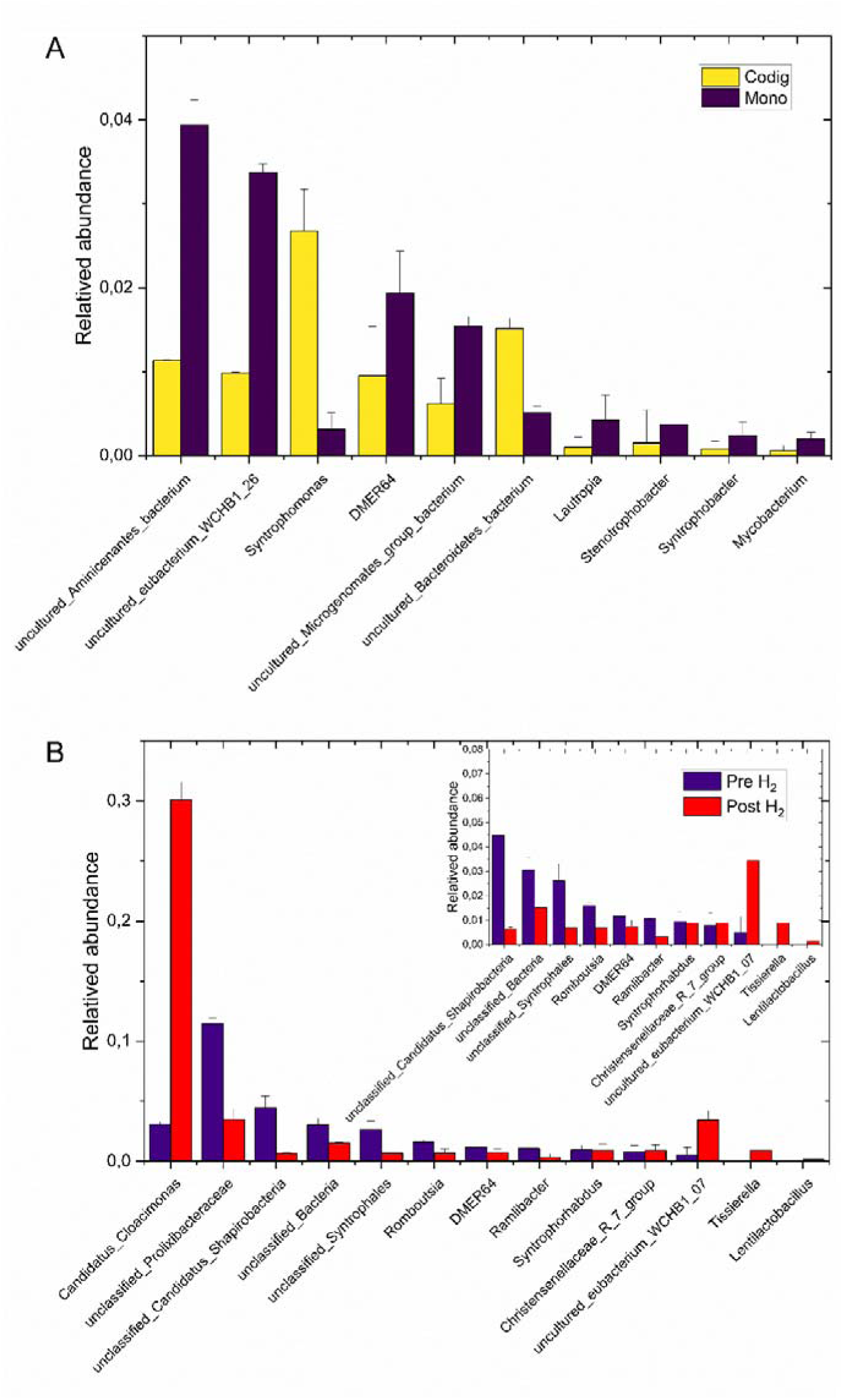
Comparative analysis of relative abundance of main bacterial Genus comparing two different conditions. A) Mono-digestion samples and Co-digestion samples; B) Hydrogen suppling samples vs non-hydrogen suppling samples.

On the other hand, when H_2_ is added to the medium, a very marked change in the bacterial population is observed. Figure 5B shows the results of the comparative study of the relative abundance of bacterial genera before and after adding H_2_. A high abundance of the genera *Cloacimonas* and *Syntrophomonas* can be observed, probably due to the synergistic effect of the cheese fats and the H_2_ added. In contrast, the relative abundance of Proteobacteria and Chloroflexi decreased under H_2_ exposure, suggesting a lower relevance of oxidative and hydrolytic routes under conditions of increased H_2_ partial pressure. The persistence of Cloacimonadota and Verrucomicrobiota was notable, supporting their role in the fermentation of intermediate compounds and the degradation of complex polysaccharides in lipid-rich environments. From a functional standpoint, the bacterial consortia under H_2_ injection showed an enrichment of taxa involved in syntrophic metabolism and interspecies electron transfer. Positive correlations were detected between *Syntrophomonas*, Ruminococcaceae, and *Methanospirillum*, the latter previously identified as the main hydrogenotrophic methanogen in Block I (Supplementary **Figure S3**). This repeated pattern in the two experimental blocks performed confirms that the co-digestion system, when supplied with external H_2_, enhances the syntrophic network established during the earlier lipid-driven adaptation phase

**Figure S5** illustrates the genus-level bacterial (A) and archaeal (B) co-occurrence networks derived from the continuous co-digestion experiments with dairy-greases wastes and during *in situ* biomethanation with H_2_. In the bacterial network **(Figure S5A)**, the addition of greases resulted in a dense cluster of positively correlated taxa associated with lipid hydrolysis, lipid degrading routes and syntrophic fatty-acid degradation. Central nodes such as *Syntrophomonas*, *Candidatus Cloacimonas*, and several members of *Firmicutes* and *Bacteroidota* showed strong positive correlations (thick orange edges), indicating tightly coordinated metabolic interactions required for the degradation of LCFAs. Its chemolithoautotrophic metabolism and its ability to grow in strictly anaerobic environments could favor complementary metabolic routes that support other fermentative microorganisms and syntrophic organisms such as Syntrophomonas, fundamental in the degradation of LCFAs and propionic acid [60]. It should be noted that cheese fats have a high concentration of propionic acid (see **Table 1**). These syntrophic bacteria depend on their partners to maintain low H_2_ partial pressure, which is essential to thermodynamically enable β-oxidation pathways. The presence of multiple negative correlations (green edges) between LCFAs-degrading specialists and taxa less adapted to lipids suggests niche differentiation and competitive exclusion as the substrate spectrum shifted towards more reduced, energy-rich grease-derived compounds. Overall, the bacterial network became more interconnected under grease co-digestion, reflecting the formation of a metabolically specialized consortium optimized for LCFAs breakdown and the production of H_2_ and acetate, which arekey intermediates for downstream methanogenesis.

The archaeal community network **(Figure S5B)** was comparably compact but dominated by strong positive correlations among hydrogenotrophic and acetoclastic methanogens. Genera such as *Methanospirillum*, *Methanolinea*, *Methanobacterium* and *Methanosaeta* formed the core of the network, linked by thick orange edges that indicate cooperative responses to the metabolic products generated by the bacterial partners. These positive associations reflect stable functional alliances required for efficient H_2_ and acetate conversion during both conventional co-digestion and *in situ* biomethanation. Notably, the introduction of H_2_ resulted in the strengthening of correlations among hydrogenotrophic groups, which increased in abundance and centrality within the network, highlighting their rapid adaptation to elevated H_2_ availability. At the same time, negative correlations (green edges) between hydrogenotrophic and some acetoclastic methanogens suggest competition for shared intermediates or shifts in substrate preference under conditions of high H_2_ dosing. A negative correlation can also be observed between *Methanospirillum* and *Methanoculleus*, both hydrogenotrophic species, probably due to competition for substrate. Minor archaeal taxa, including members of *Thermoplasmatota*, maintained weaker but persistent connections, indicating auxiliary roles in carbon turnover or in stabilizing redox conditions within the digester.

Together, the bacterial and archaeal networks demonstrate that dairy greases co-digestion promotes the emergence of a syntrophic bacterial consortium capable of efficiently degrading LCFAs, while H_2_ addition reinforces a hydrogenotrophic methanogenic community highly adapted to *in situ* biomethanation under mesophilic conditions. The strong and coherent correlations across both domains highlight the establishment of an integrated microbial network that supports enhanced CH_4_ formation under grease-rich-substrate and hydrogen-enriched conditions.

Figure 6 summarizes how process and environmental conditions and reactor performance are related to bacterial (Fig. 6A) and archaeal (Fig. 6B) community structure during co-digestion with dairy greases and *in situ* biomethanation. In the heatmap, most operational and performance variables co-varied in an expected manner.. Importantly, externally added H_2_ (H_2_ added) was positively correlated with CH_4_ content and biogas production and tended to show an inverse relationship with CO_2_, reflecting the progressive conversion of CO_2_ to CH_4_ during the biomethanation stages with H_2_ dosing.

**Figure 6.**
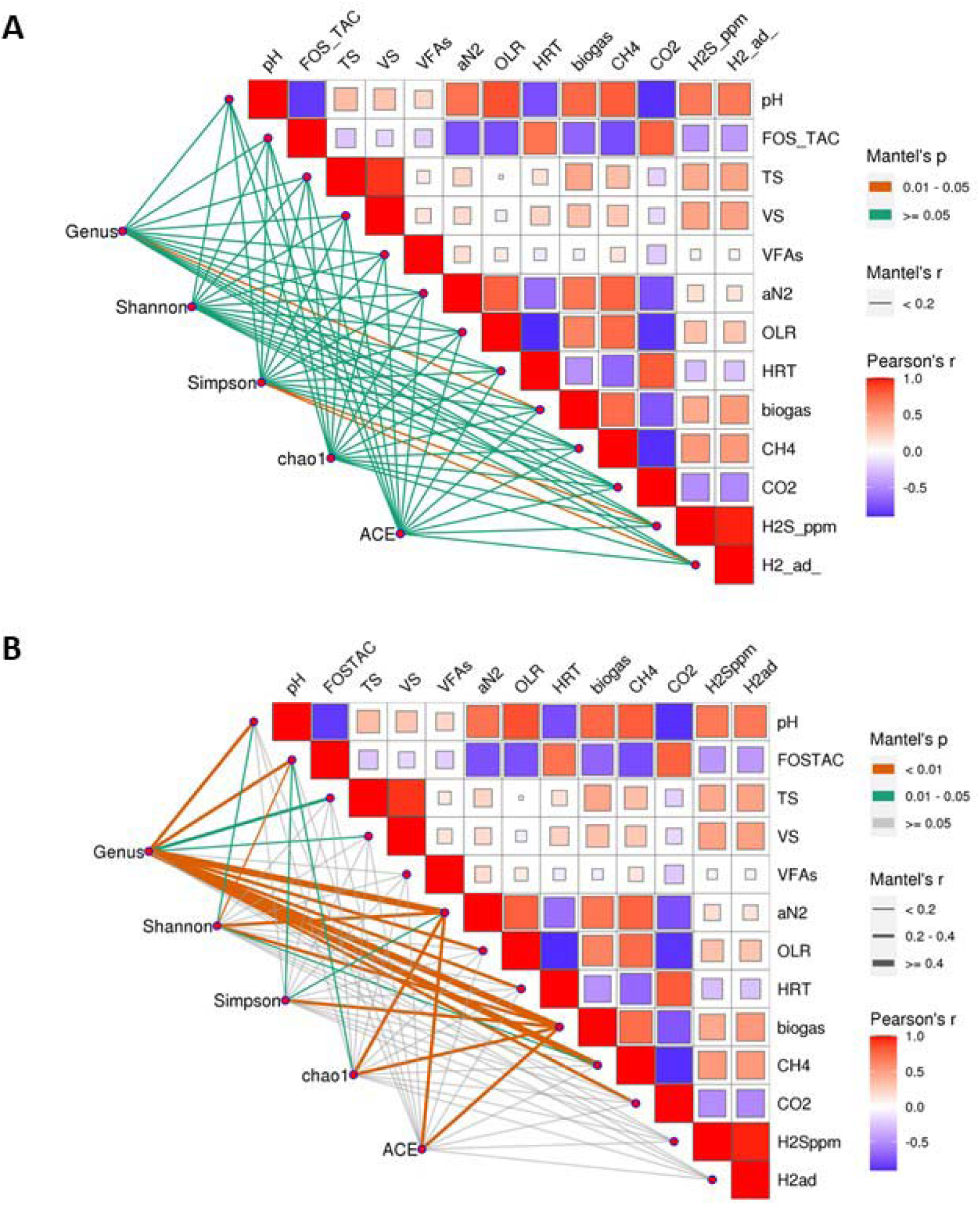
Combined heatmap–network diagrams showing the correlations between environmental parameters, alpha-diversity indices, and microbial taxa for. (A) bacteria and (B) archaea. Environmental parameters include pH, FOS/TAC ratio, total solids (TS), volatile solids (VS), volatile fatty acids (VFAs), ammoniacal nitrogen (AN_2_), organic loading rate (OLR), hydraulic retention time (HRT), biogas production, methane (CH4), carbon dioxide (CO_2_), hydrogen sulfide (_H2S_), and externally supplied hydrogen (H_2_ added). The heatmap (top right) displays positive and negative Pearson correlations among environmental factors. The network (bottom left) depicts relationships between microbial taxa, diversity indices, and environmental variables, where line color represents Mantel p-values and line thickness corresponds to Mantel r-values,

The bacterial network (Figure 6A) shows that bacterial alpha-diversity is significantly structured by these environmental gradients. The network in the lower-left panel depicts Mantel correlations between community indices and environmental variables. Most connections correspond to non-significant associations (green edges, p ≥ 0.05), indicating that several operational gradients influence bacterial diversity but with limited statistical support. In contrast, significant Mantel correlations (orange edges, p = 0.01–0.05) are primarily associated with biogas production, H_2_S production, and H_2_ addition (H_2__ad). These significant links reveal that gas-related parameters exert the strongest measurable effects on bacterial community structure. Specifically, significant positive associations connecting Shannon, Simpson, and genus richness with biogas and H_2__ad suggest that richer and more even bacterial assemblages developed when H_2_ supplementation stimulated metabolic interactions and gas formation. Significant correlations with H_2_S indicate shifts in sulfur-transforming taxa under these conditions. Conversely, the richness estimators Chao1 and ACE exhibit few or no significant edges, supporting the interpretation that changes in evenness and dominance (Shannon and Simpson), rather than total estimated richness, better capture the bacterial community response to operational variation.

The archaeal community network (Figure 6B) is characterized by fewer but generally stronger and more significant associations (numerous thick orange edges, p < 0.01) between diversity indices and process variables. Genus richness and Shannon/Simpson diversity of archaea were tightly linked to OLR, biogas production and CH_4_ content, whereas relationships with solids and VFAs were comparatively weaker. This suggests that methanogenic community structure is governed primarily by the energy yield and electron-donor availability, rather than by bulk organic matter concentrations. As in the bacterial network, Chao1 and ACE contributed little to explaining archaeal responses, which implies that changes in the relative dominance of key methanogenic genera are more relevant than the appearance of many new low-abundance taxa.

A functional overview of the bacterial community using FAPROTAX (Supplementary Figure S4) further supports the enrichment of hydrolytic–fermentative populations under co-digestion with dairy greases. In both operational modes, the dominant predicted function was “fermentation”, but its contribution was clearly higher in the co-digestion samples, where it accounted for roughly twice the mean proportion observed during sludge mono-digestion, with a significant difference between groups (corrected *p* < 0.05). This enrichment of fermentation-related functions is consistent with the selection of hydrolytic and fermentative bacteria capable of depolymerizing complex organic matter and converting lipids and proteins from cheese-waste streams into VFAs, H_2_ and acetate, in line with the syntrophic LCFAs degraders highlighted in the community networks (Figure S5). In contrast, other predicted functions (e.g., nitrogen transformations) remained at very low relative abundances and showed only minor differences between treatments, indicating that the main functional shift induced by grease co-digestion was the reinforcement of the primary hydrolytic–fermentative metabolism that fuels downstream.

## 4. Conclusions

Co-digestion of municipal wastewater sludge with dairy industry greases profoundly reshaped the structure and function of the anaerobic microbiome. Lipid addition shifted the community from a Chloroflexi–Proteobacteria background with *Methanosaeta*– and *Methanofastidiosum*-dominated methanogenesis towards Firmicutes-, Bacteroidota-, Cloacimonadota– and Verrucomicrobiota-rich consortia coupled to hydrogenotrophic archaea, particularly *Methanospirillum* and *Methanosaeta*. This compositional shift redirected methanogenesis from predominantly acetoclastic to hydrogenotrophic pathways, strengthening syntrophic links between β-oxidizing / lipolytic bacteria and hydrogenotrophic methanogens. The consistent enrichment of *Methanospirillum* suggests its potential use as a bioindicator of well-performing, hydrogen-enriched digesters treating lipid-rich residues.

Stepwise increases in OLR combined with external H_2_ addition demonstrated that mesophilic digesters co-fed with dairy greases can sustain high CH_4_ yields while simultaneously upgrading biogas to 80 – 82 % CH_4_. Injected H_2_ was efficiently converted (≈70 – 90% up to 40% of the stoichiometric requirement), confirming that the supplied H_2_ was largely channeled to microbial CO_2_ reduction rather than lost to the gas phase. H_2__ad reinforced the syntrophic network previously selected by lipid co-digestion, enabling rapid H_2_ consumption and stable CH_4_ formation. Multivariate and network analyses showed that bacterial diversity closely followed gradients in loading, solids and acidification, whereas archaeal diversity was tightly coupled to gas-phase variables and H_2_ addition, consistent with a community dominated by hydrogenotrophic methanogens.

Overall, co-digestion with dairy greases followed by intermittent H_2_ dosing produced a resilient mesophilic consortium able to couple β-oxidation-derived electron flow with hydrogenotrophic methanogenesis. The novelty of this work lies not in the individual use of lipid co-digestion or mesophilic in situ biomethanation, both of which have been previously described, but in demonstrating that lipid-driven microbial selection can be intentionally harnessed to establish a robust hydrogenotrophic consortium capable of sustaining efficient biomethanation under mesophilic conditions. By coupling these two concepts, conventional anaerobic digesters can be transformed into effective H_2_-utilization platforms without thermophilic operation, pressurization, gas recirculation, or substantial process modifications, thereby providing a realistic pathway for integrating renewable hydrogen into existing wastewater treatment infrastructure.

## Appendix A. Supplementary data

The following are the Supplementary data to this article:

## Author Approvals

All authors have seen and approved the manuscript.

## Acknowledgements

This research was carried out in line with the BIOUP and BIOFORAIR research project. Project BIOUP, CPP2021-009086, is one of the 2021 public-private collaborative projects funded by MCIN/AEI/10.13039/501100011033 and the European Union – Next GenerationEU/PRTR. National Project BIOFORAIR, which has received funding from the Ministerio de Ciencia Innovacion y Universidades – Generación Conocimiento Program under grant agreement n°. <u>PID2022–1386820B-I00</u>. A. Lao-Zea would like to acknowledge the financial support of this Ministry (<u>PTA2022–022087-I</u>). The authors gratefully acknowledge Entrepinares S.A.U. for providing the dairy wastewater grease used as co-substrate in this study.

## CRediT authorship contribution statement

**MLR**: Conceptualization; Methodology; Supervision; Project administration; Investigation; Data curation; Formal analysis; Visualization. **ALZ**: investigation; Writing – original draft; Writing – review and editing. **ADM:** Formal analysis; Writing – original draft; Writing – review and editing. **FF**: Formal analysis; Writing – original draft; Writing – review and editing. **ID:** methodology; Writing – review and editing**. JCG**: Investigation. **RI**: Resources. **SS:** Methodology, Data Curation; Writing – review and editing. **MGA**: Conceptualization; Methodology; Supervision; Project administration; Investigation; Data curation; Formal analysis; Visualization; Writing – original draft; Writing – review and editing.

## Data availability

Data will be made available on request.

## Nomenclature

AD: anaerobic digestion
CH_4_: methane
CO_2_: carbon dioxide
COD: chemical oxygen demand
DG: dairy grease
GHG: greenhouse gas
H_2_: Hydrogen
H_2__ad: Hydrogen addition
HPLC: high-performance liquid chromatography
LCFAs: long-chain fatty acids
NH_4_-N: ammoniacal nitrogen
PO_4_–P: orthophosphate
SL: sludge
Ss: sewage sludge
TIC: total inorganic carbon
TKN: total Kjeldahl nitrogen
TOC: total organic carbon
TP: total phosphorous
TS: total solids
VFAs: volatile fatty acids
VS: volatile solids
WWTP: wastewater treatment plant

