## Appendix A for "Dairy wastewater grease stabilizes *in situ* mesophilic biomethanation for H_2_-to-CH_4_ conversion"

^1^Advanced Biofuels and Bioproducts Unit, Department of Energy. Centro de Investigaciones Energéticas, Medioambientales, and Tecnológicas (CIEMAT). Madrid. Spain

^2^FOTOAIR Unit, Department of Energy. Centro de Investigaciones Energéticas, Medioambientales, and Tecnológicas (CIEMAT). Madrid. Spain

^3^Innovation Department, Acciona Agua S.A.U. Avda. de les Garrigues, 22, 08820 El Prat de Llobregat, Barcelona, Spain

^4^Institute of Sustainable Process, University of Valladolid (UVa), 47011, Valladolid, Spain

^5^Department of Chemical Engineering and Environmental Technology, University of

Valladolid, Paseo Prado de la Magdalena 3-5, 47011 Valladolid, Spain

Correspondence to: Miguel G. Acedos. Advanced Biofuels and Bioproducts Unit, Department of Energy. Centro de Investigaciones Energéticas, Medioambientales, and Tecnológicas (CIEMAT). Avenida Complutense 40, 28040 Madrid. Spain

Phone number M.G. Acedos: +34 914102692

**Tables and figures**

**Table S1.** Primers for bacterial and archaea sequencing

| Primer | Sequence |
| --- | --- |
| Bacteria 16S rRNA F | 5'- AGRGTTTGATYNTGGCTCAG-3' |
| Bacteria 16S rRNA R | 5'-TASGGHTACCTTGTTASGACTT-3' |
| Archeae 16S rRNA F | 5'-GKTTGATCCYGSCRGAG -3 |
| Archeae 16S rRNA R | 5'-GGYTACCTTGTTACGACTT-3' |

**Table S2**. Nomenclature and list of samples analysed by metagenomics for the preliminary experiments. OLR (organic loading rate in g_VS_.L^-1^.d^-1^)

| Sample Nomenclature | Experimental Sample |
| --- | --- |
| SC1 | R1 time 0 days (OLR=0.5) |
| SC2 | R2 time 0 days (OLR=0.5) |
| SC3 | R3 time 0 days (OLR=0.5) |
| SC4 | R1 time 40 days (OLR=1) |
| SC5 | R2 time 40 days (OLR=1) |
| SC6 | R3 time 40 days (OLR=1) |
| SC7 | R1 time 70 days (OLR=1.25) |
| SC8 | R1. Time 90 days (end) (OLR=1.5) |
| SC9 | R2 time 170 days (end) (OLR=1.5) |
| SC10 | R3 time 170 days (end) (OLR=1.5) |

**Table S3.** Summary of average specific production rates and composition of biogas produced for experiment R4 of the second experimental block

| Scenario | Biogas Production  (Nml_biogas._L^-1^.d^-1^) | CH_4_  (%) | CO_2_  (%) | H_2_S  (ppm) | H_2_  (%) |
| --- | --- | --- | --- | --- | --- |
| I | 374 ± 52 | 52.6 ± 2.6 | 47.4 ± 2.5 | 3 ± 1 | - |
| II | 839 ± 54 | 63.1 ± 0.9 | 36.2 ± 0.3 | 5 ± 2 | - |
| III | 1225 ± 120 | 67.2 ± 1.1 | 33.2 ± 1.6 | 19 ± 1 | - |
| a | 1123 ± 244 | 70.5 ± 1.0 | 29.5 ± 0.9 | 108 ± 36 | 0.4 ± 0.1 |
| b | 1068 ± 128 | 70.4 ± 1.5 | 29.0 ± 1.8 | 152 ± 19 | 0.2 ± 0.1 |
| c | 1180 ± 145 | 71.5 ± 1.5 | 27.7 ± 2.3 | 156 ± 29 | 0.8 ± 0.6 |
| d | 1279 ± 198 | 74.1 ± 1.7 | 25.8 ± 1.5 | 291 ± 54 | 3.2 ± 1.4 |
| e | 1327 ± 210 | 72.6 ± 4.9 | 26.5 ± 4.3 | 500 ± 126 | 13.0 ± 3.0 |
| f | 1702 ± 406 | 79.8 ± 2.3 | 21.8 ± 3.8 | 336 ± 39 | 5.3 ± 1.3 |

**Table S4.** Nomenclature and list of samples analysed by metagenomics for the second experimental block. R11-R14 refer to reactor 1 (R1), monodigestion of sewage sludge. R21-R29 refer to reactor 2 and R31-R39 refer to reactor 3 (R2 and R3), co-digestion of sewage sludge and dairy WWTP greases, without hydrogen supply; and R41-R411 and its duplicates R41d-R411d, refer to reactor 4 (R4), co-digestion of sewage sludge and dairy WWTP greases, and hydrogen supply from sample R45/R45d to R411/R411d.

| Sample Nomenclature | Experimental Sample for Metagenomic Sequencing |
| --- | --- |
| R11 | R1 time 0 days |
| R12 | R1 end of OLR I |
| R13 | R1 end of OLR II |
| R14 | R1 end of OLR III (trial end) |
| R21 | R2 time 0 days |
| R22 | R2 end of OLR I |
| R23 | R2 end of OLR II |
| R24 | R2 OLR III time 1 |
| R25 | R2 OLR III time 2 |
| R26 | R2 OLR III time 3 |
| R27 | R2 OLR III time 4 |
| R28 | R2 OLR III time 5 |
| R29 | R2 OLR III time 6 (assay end) |
| R31 | R3 time 0 days |
| R32 | R3 end of OLR I |
| R33 | R3 end of OLR II |
| R34 | R3 OLR III time 1 |
| R35 | R3 OLR III time 2 |
| R36 | R3 OLR III time 3 |
| R37 | R3 OLR III time 4 |
| R38 | R3 OLR III time 5 |
| R39 | R3 OLR III time 6 (assay end) |
| R41/R41d | R4 time 0 days |
| R42/R42d | R4 end of OLR I |
| R43/R43d | R4 end of OLR II |
| R44/R44d | R4 OLR III 0 % H_2_ |
| R45/R45d | R4 OLR III 5 % H_2_ |
| R46/R46d | R4 OLR III 10 % H_2_ |
| R47/R47d | R4 OLR III 20 % H_2_ |
| R48/R48d | R4 OLR III 40 % H_2_ |
| R49/R49d | R4 OLR III 60 % H_2_ |
| R410/R410d | R4 OLR III 50 % H_2_ |
| R411/R411d | R4 OLR III 50 % H_2_ (assay end) |

**Figures**

| 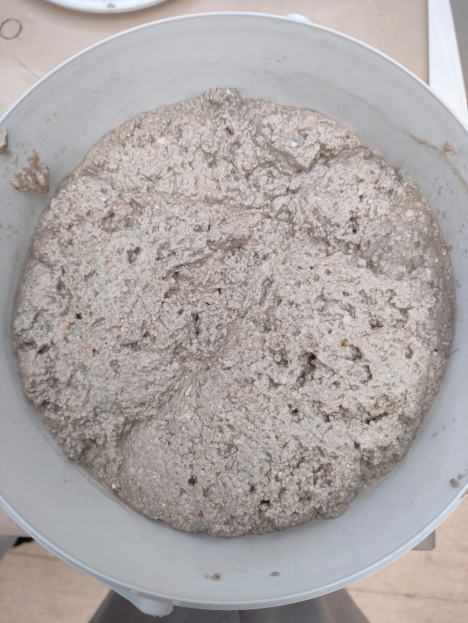 | 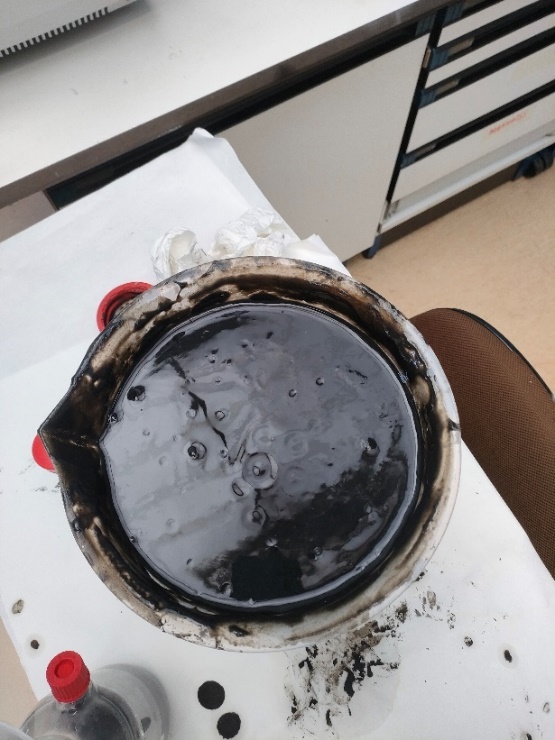 | 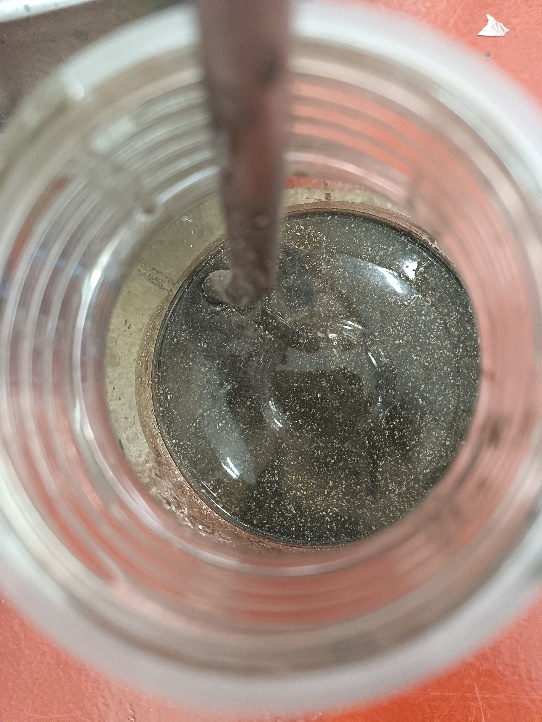 |
| --- | --- | --- |
| Dairy Greases | Sewage sludge | Co-digestion substrate |

**Figure S1.** Photograph of the substrates used in this study. From left to right: dairy greases municipal sewage sludge; co-digestion substrate

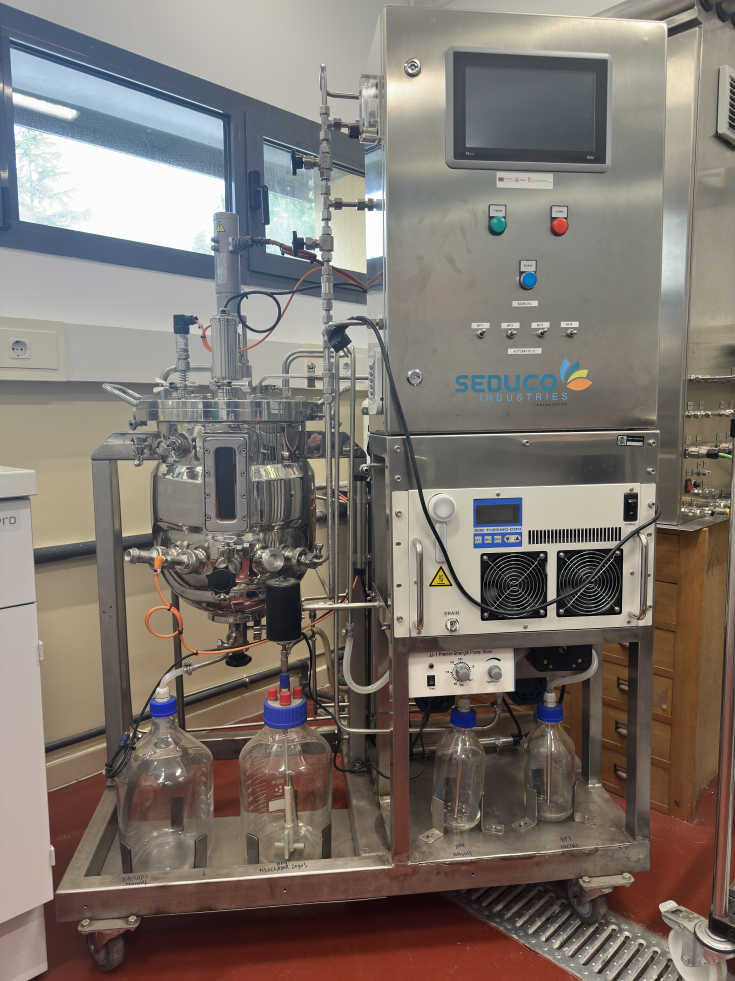

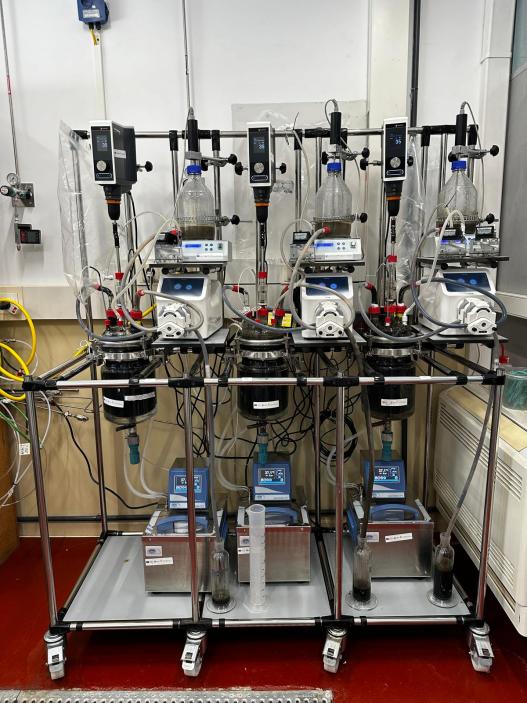

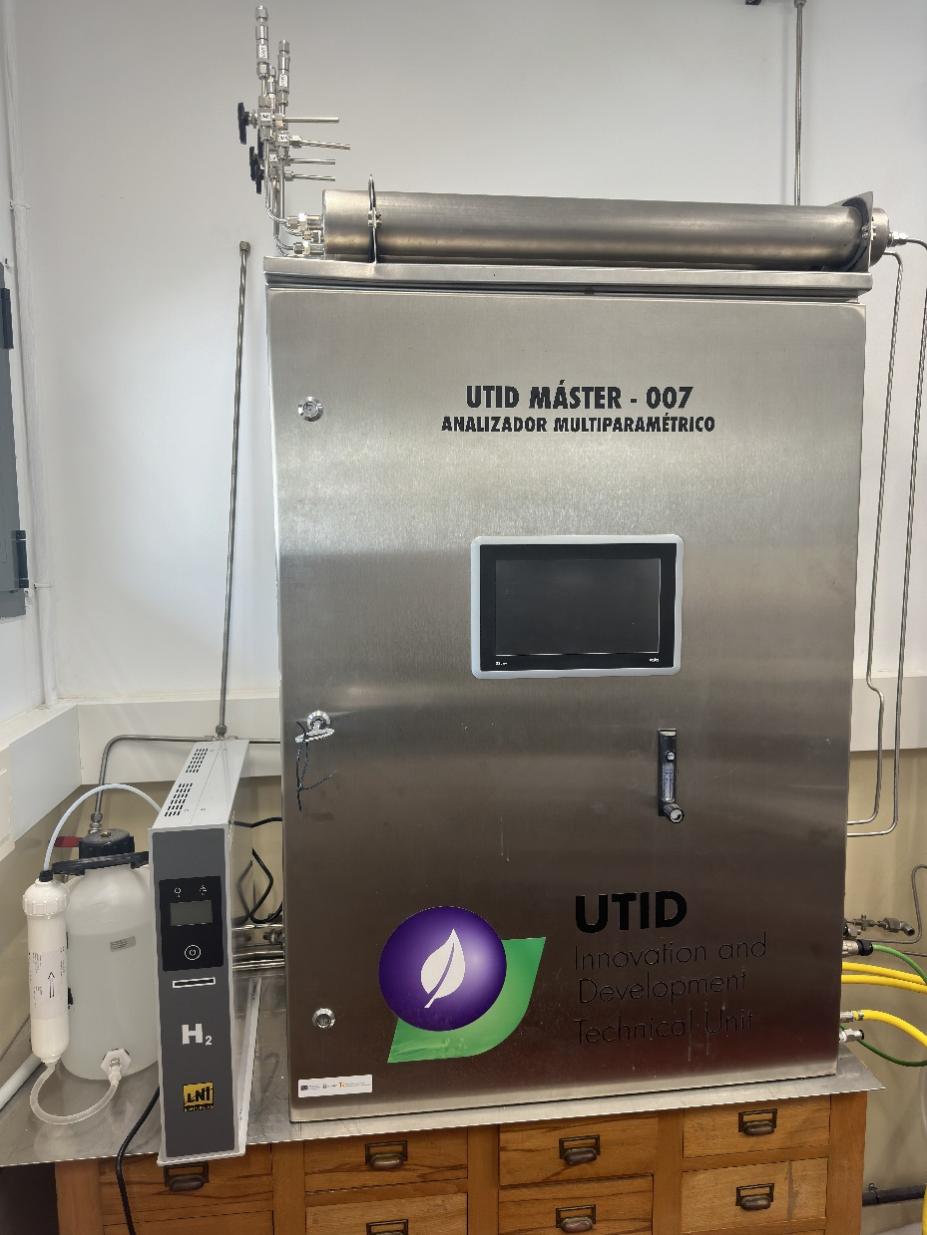

**Figure S2.** 20 L digester, 3 L digesters and electrolyser employed in this study.

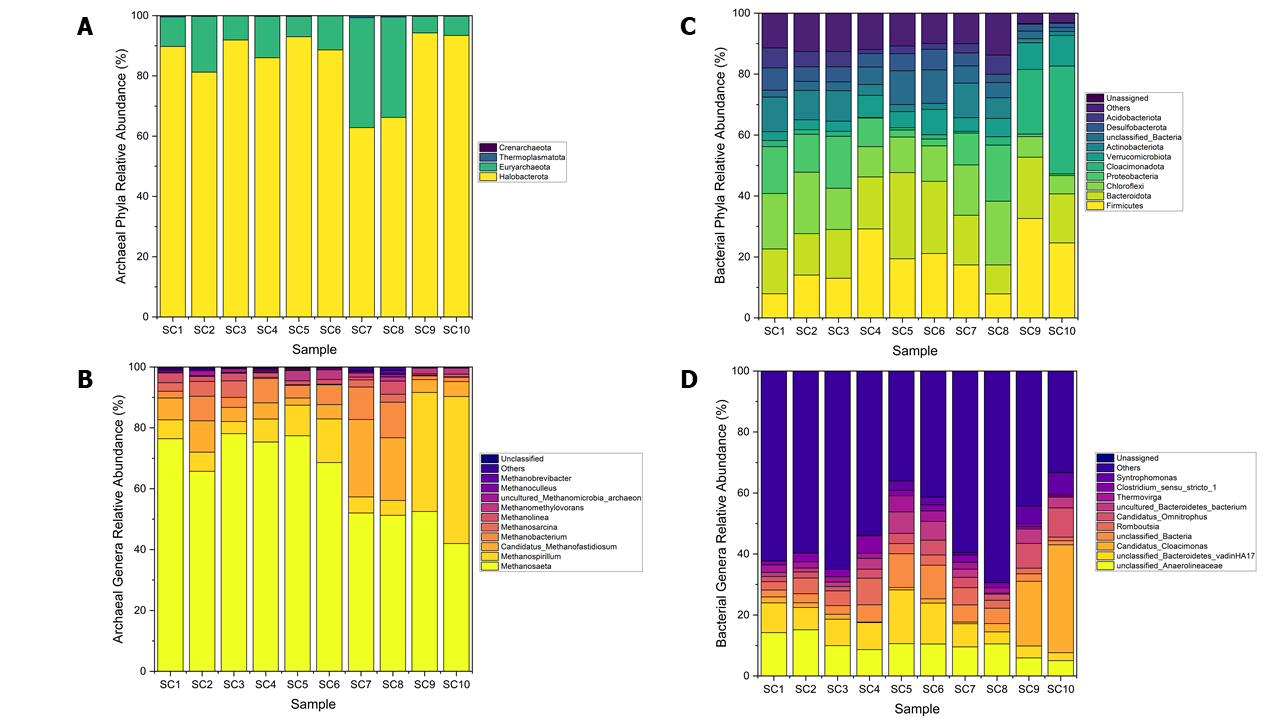

**Figure S3.** Relative abundance of archaeal phyla (A) and archaeal genera (B) in the samples analysed, and relative abundance of bacterial phyla (C) and bacterial genera (D) in the samples analysed, for the preliminary experiments.

In the preliminary experiments. Reactors R1-R3 were fed with either pretreated sewage sludge alone (R1) or a co-digestion mixture of pretreated sludge and dairy greases (80:20 w/w; R2 and R3). Biogas production increased with OLR in all cases, but the co-digestion reactors consistently showed higher methane yields than the sludge mono-digestion. Maximum methane productivity reached approximately 0.45 L CH_4_ g_VS_ ⁻¹ at OLR 2.5 g_VS._L⁻1d⁻¹, confirming the positive effect of lipid-rich residues.

In this preliminary experiments, which was carried out under conditions similar to those presented in the manuscript, the same behavior of the microbial population was observed. On the one hand, an enrichment in hydrogenotrophic archaea was observed when cheese fats and lipolytic bacteria that favor the development of hydrogenotrophic archaea were used. The samples are described in the table S3.

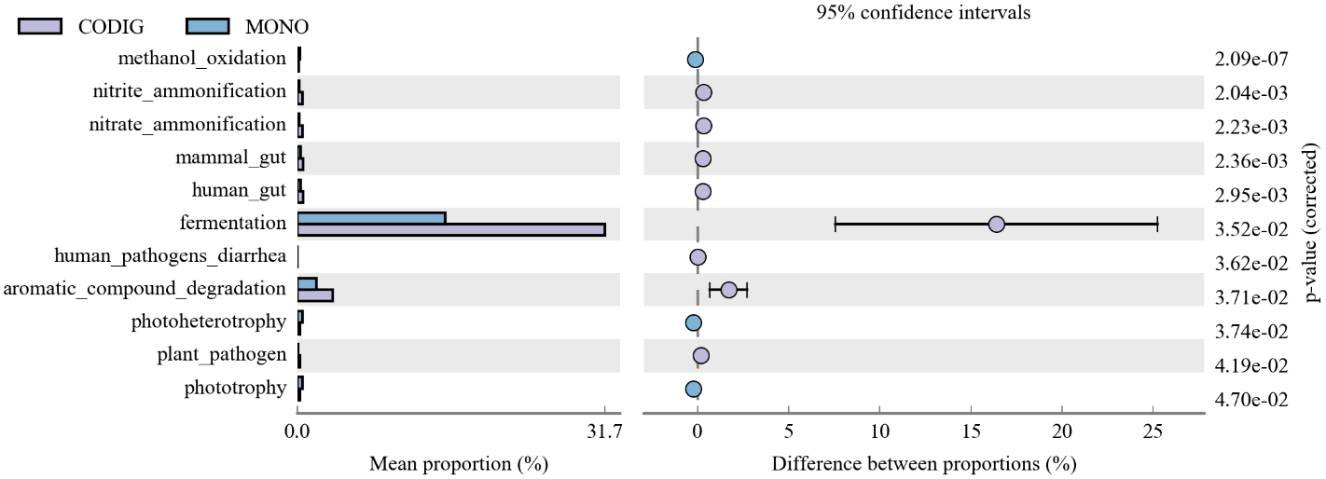

**Figure S4.** Analysis of FAPROTAX difference between groups. Comparative analysis between co-digestion samples and monodigestion samples for bacterial community.

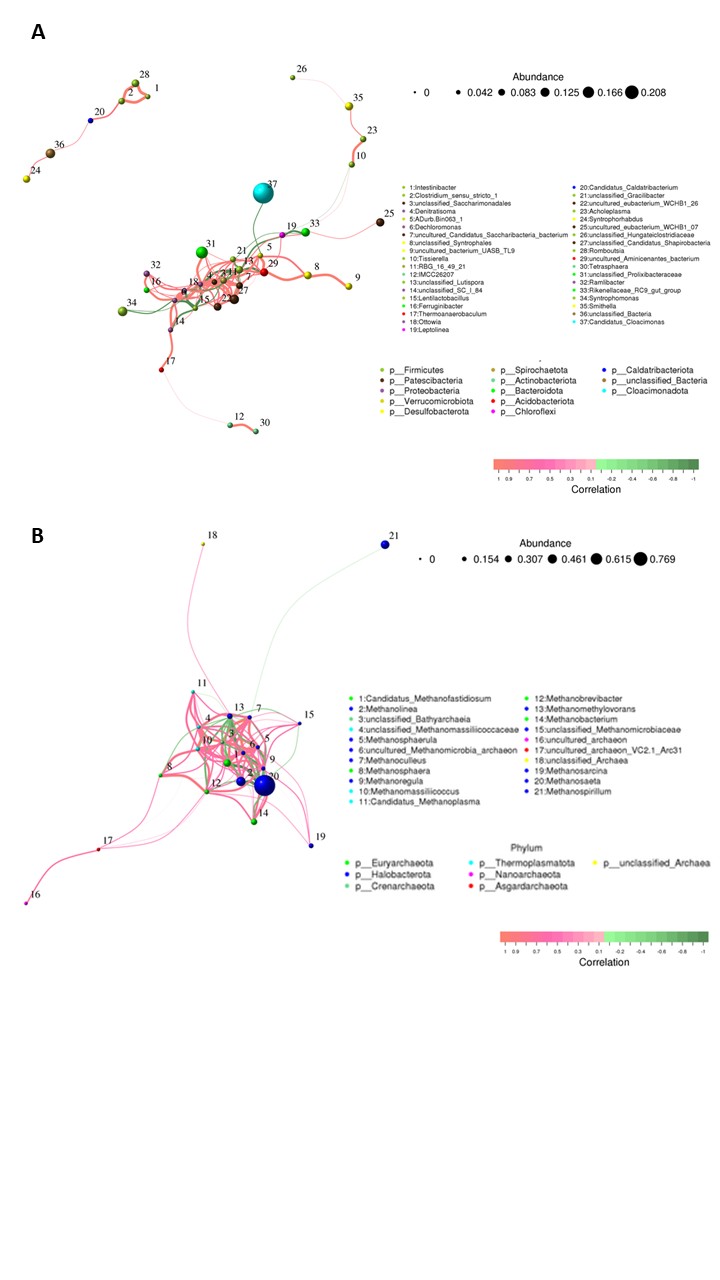

**Figure S5.** Bacterial Species network at genus level (A) and archaea (B) species network at genus level. Circles represent species, size of circle represents the abundance; the edges represent the correlation between the two species, the thickness of the edge represents the strength of the correlation, and the color of the line: orange represents the positive correlation, while green represents the negative correlation.

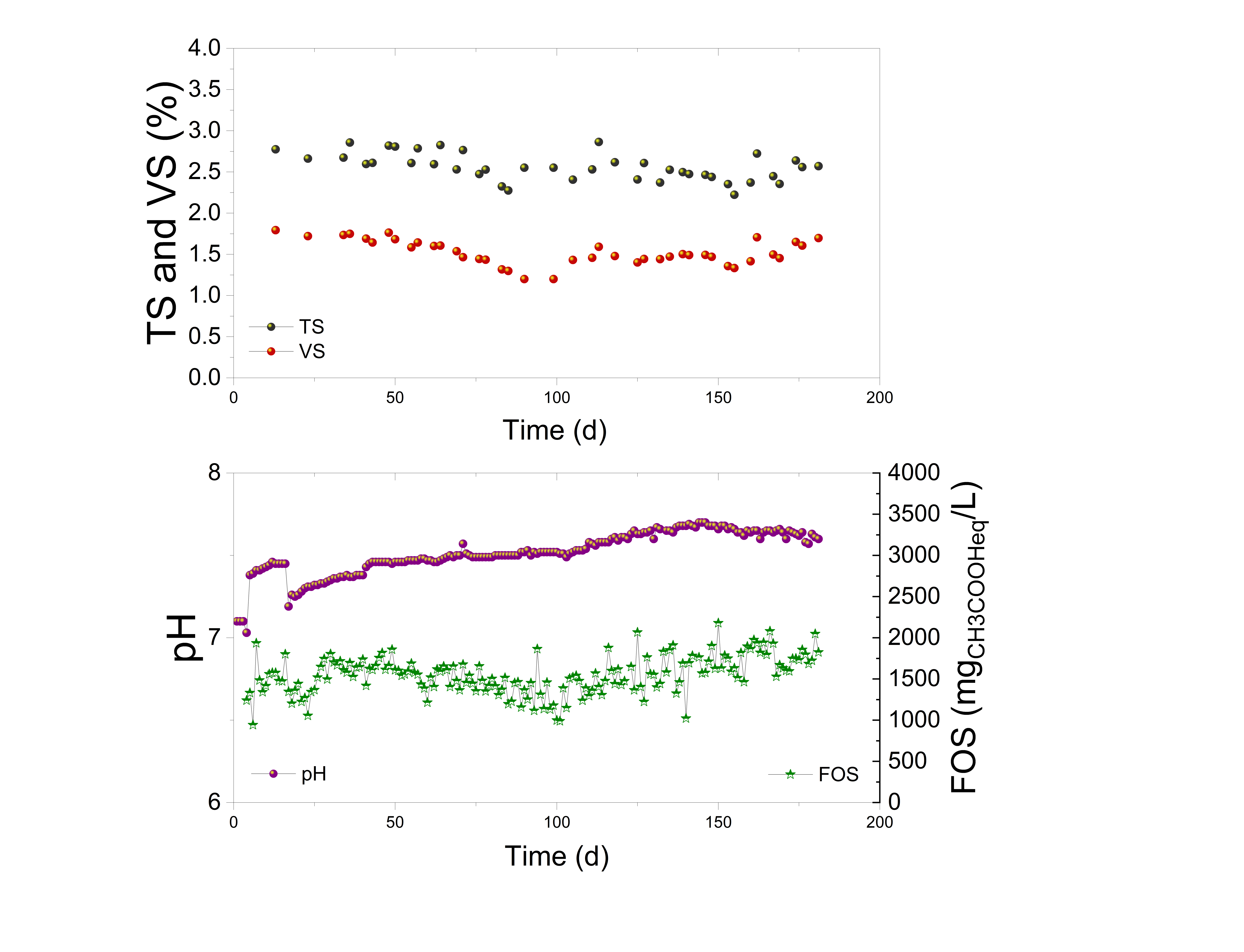

**Figure S6.** Evolution of total solids (TS) and volatile solids (VS) (top panel), and pH (left axis) and FOS (right axis, expressed as mg acetic acid equivalents L⁻¹) (bottom panel) in reactor R1. Reactor configuration and operational conditions correspond to those described for R1 in Table 2 and Figures 1 and 2 of the main manuscript. This figure specifically refers to the reactors presented in the main study to avoid confusion with additional experiments included in the Supplementary Information.

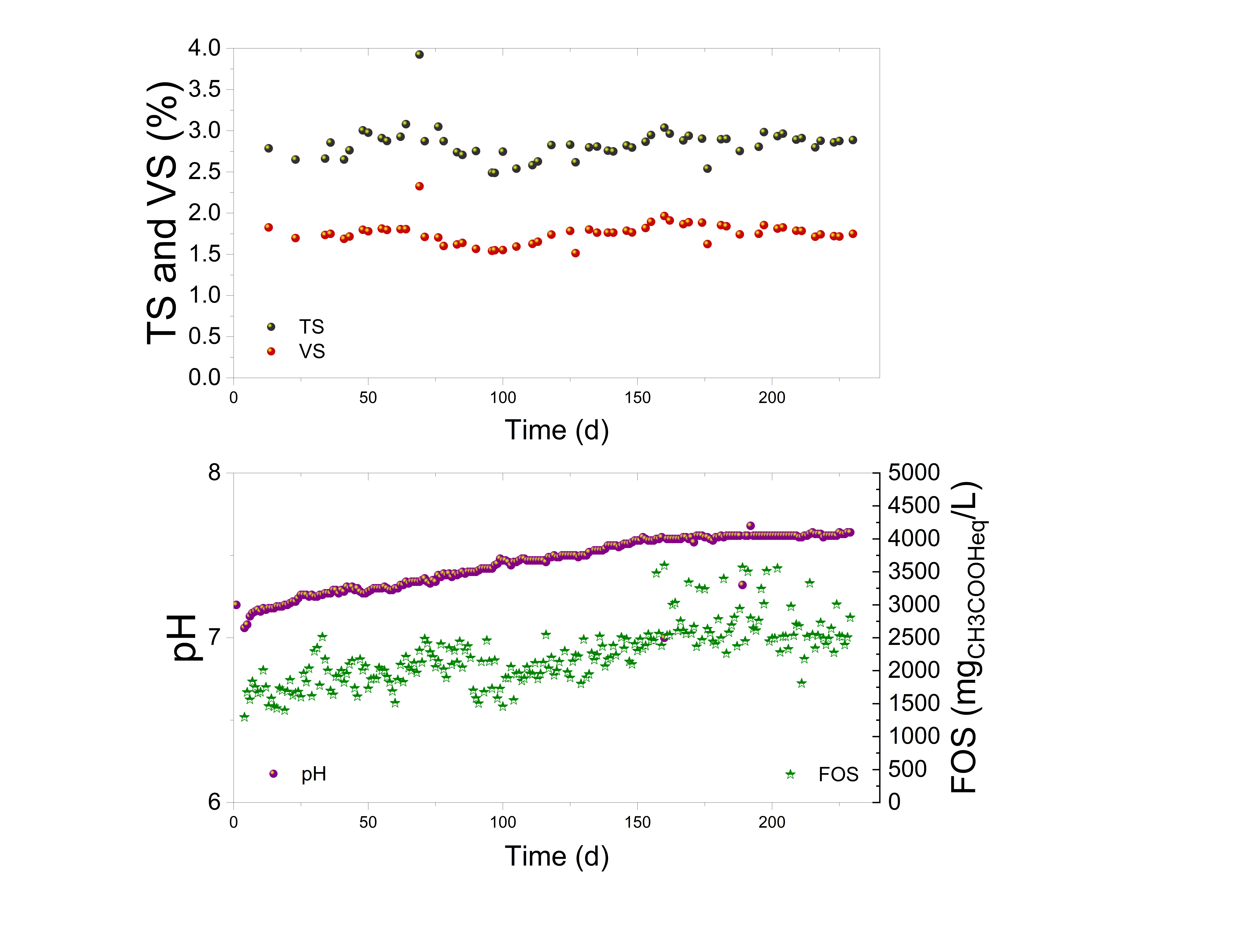

**Figure S7.** Evolution of total solids (TS) and volatile solids (VS) (top panel), and pH (left axis) and FOS (right axis, expressed as mg acetic acid equivalents L⁻¹) (bottom panel) in reactor R2. Reactor configuration and operational conditions correspond to those described for R2 in Table 2 and Figures 1 and 2 of the main manuscript. This figure specifically refers to the reactors presented in the main study to avoid confusion with additional experiments included in the Supplementary Information.

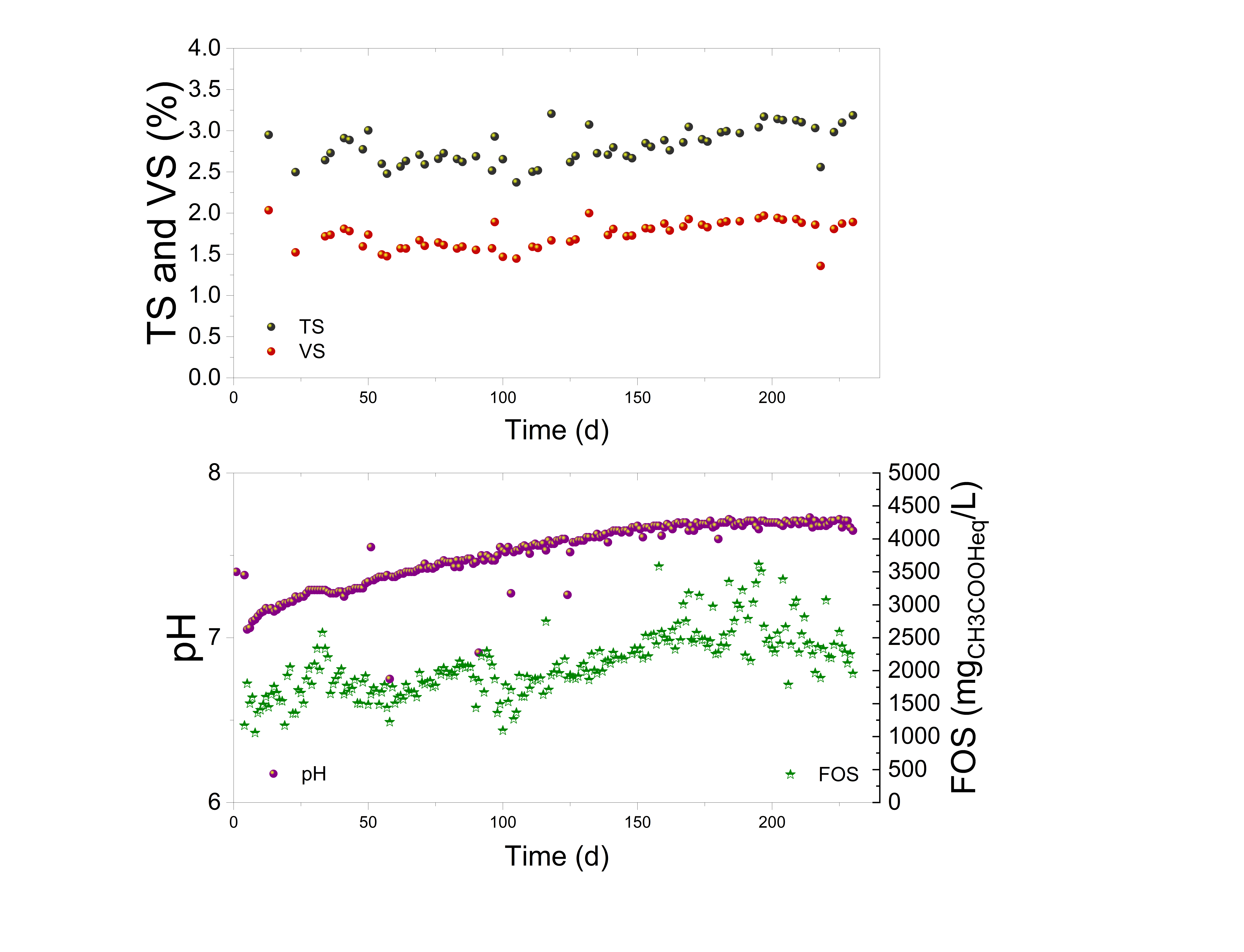

**Figure S8.** Evolution of total solids (TS) and volatile solids (VS) (top panel), and pH (left axis) and FOS (right axis, expressed as mg acetic acid equivalents L⁻¹) (bottom panel) in reactor R3. Reactor configuration and operational conditions correspond to those described for R3 in Table 2 and Figures 1 and 2 of the main manuscript. This figure specifically refers to the reactors presented in the main study to avoid confusion with additional experiments included in the Supplementary Information.

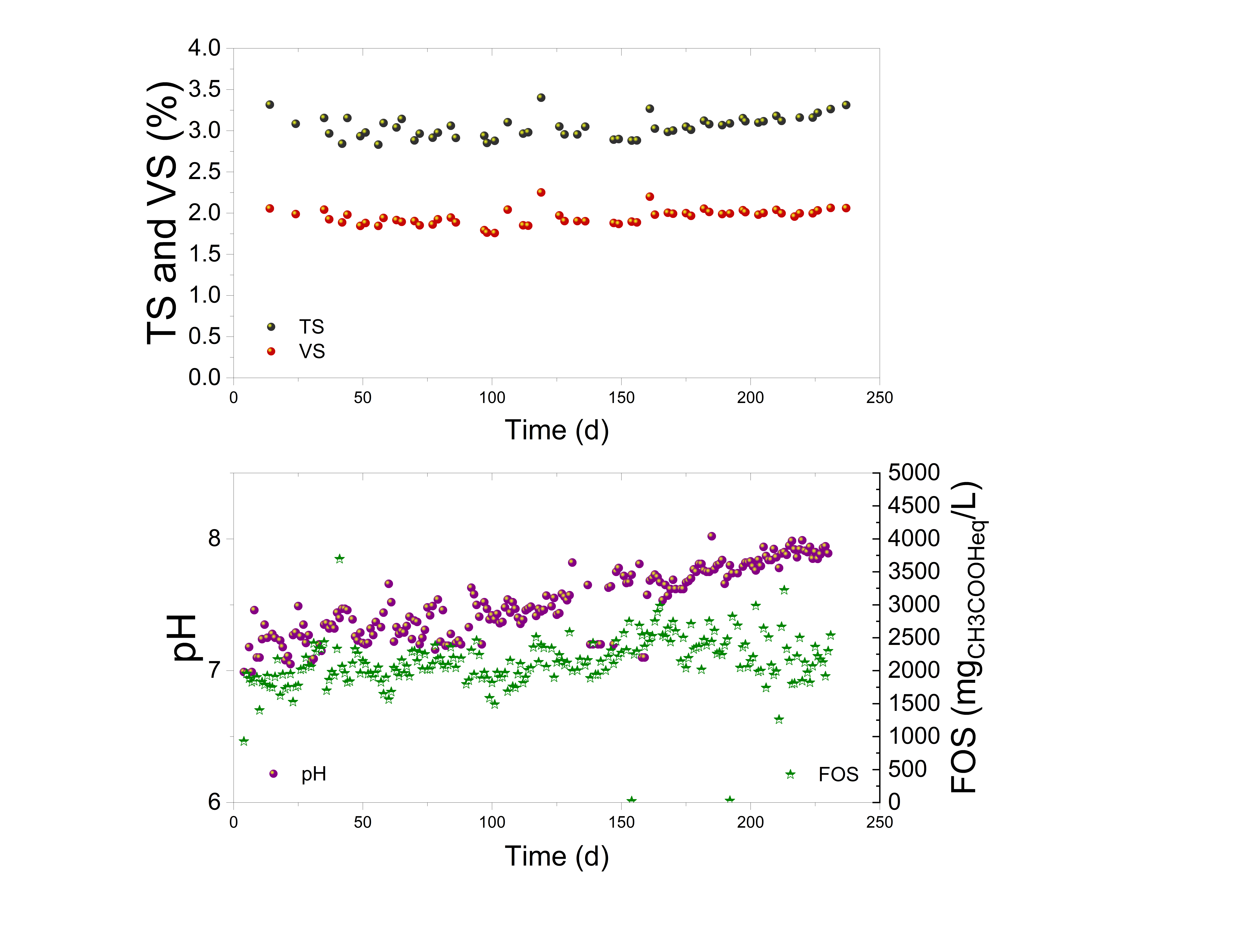

**Figure S9.** Evolution of total solids (TS) and volatile solids (VS) (top panel), and pH (left axis) and FOS (right axis, expressed as mg acetic acid equivalents L⁻¹) (bottom panel) in reactor R4. Reactor configuration and operational conditions correspond to those described for R4 in Table 2 and Figures 1 and 2 of the main manuscript. This figure specifically refers to the reactors presented in the main study to avoid confusion with additional experiments included in the Supplementary Information.
